# Proteomics of protein trafficking by *in vivo* tissue-specific labeling

**DOI:** 10.1101/2020.04.15.039933

**Authors:** Ilia A. Droujinine, Dan Wang, Yanhui Hu, Namrata D. Udeshi, Luye Mu, Tanya Svinkina, Rebecca Zeng, Tess Branon, Areya Tabatabai, Justin A. Bosch, John M. Asara, Alice Y. Ting, Steven A. Carr, Norbert Perrimon

## Abstract

Secreted interorgan communication factors encode key regulators of homeostasis. However, long-standing questions surround their origins/destinations, mechanisms of interactions, and the number of proteins involved. Progress has been hindered by the lack of methodologies for these factors’ large-scale identification and characterization, as conventional approaches cannot identify low-abundance factors and the origins and destinations of secreted proteins. We established an *in vivo* platform to investigate secreted protein trafficking between organs proteome-wide, whereby engineered promiscuous biotin ligase BirA*G3 (a relative of TurboID) biotinylates all proteins in a subcellular compartment of one tissue, and biotinylated proteins are affinity-enriched and identified from distal organs using quantitative mass spectrometry. Using this platform, we identified 51 putative muscle-secreted proteins from heads and 269 fat body-secreted proteins from legs/muscles, of which 60-70% have human orthologs. We demonstrate, in particular, that conserved fat body-derived novel interorgan communication factors CG31326, CG2145, and CG4332 promote muscle activity. Our results indicate that the communication network of secreted proteins is vast, and we identified systemic functions for a number of these factors. This approach is widely applicable to studies in interorgan, local and intracellular protein trafficking networks, non-conventional secretion, and to mammalian systems, under healthy or diseased states.

**One Sentence Summary:** We developed an *in vivo* platform to investigate protein trafficking between organs proteome-wide, provide a resource for interorgan communication factors, and determined conserved adipokines that affect muscles.

## Main Text

Local tissue homeostasis is becoming increasingly well-understood. However, the physiological importance and presence of secreted interorgan communication factors is only beginning to be documented from experiments in *Drosophila* and vertebrates. Secreted factors acting directly or indirectly between organs encode key regulators of systemic homeostasis (*1*). These factors traffic, or translocate, intracellularly from their production sites within cells (*2*) to distal organs through blood (*1*). For instance, adipokines including leptin and adiponectin encode adipose tissue-derived systemic metabolic regulators (*1*). In addition, myokines such as irisin (cleaved form of FNDC5) and interleukin-6 are secreted by muscles to control metabolism in adipose tissue (*1*). Despite their importance, identification of interorgan communication factors is technically challenging, and a number of published results were later determined to be irreproducible or controversial (*1, 3—5*). Also, origins and/or destinations of factors including glucagon-like peptide 1 (GLP-1), ghrelin, leptin, cholecystokinin (CCK), and growth differentiation factor 11 (GDF-11) need to be clarified (*1, 5*). Moreover, because large-scale screening methods are lacking, the long-standing biological question of how many proteins are transferred between any two given organs is yet to be addressed (*1*). While a number of factors have been recognized in mammals, many physiologically-important factors likely remain to be identified (*1*).

A powerful approach to identify secreted factors from the blood is targeted mass spectrometry (MS). While MS is a sensitive method, unprocessed blood samples are exceedingly complex with large dynamic ranges of protein concentrations – the majority of the protein mass consists of a small number of protein species which dominate the MS signal, making identification of lower-abundance proteins challenging (*6*). Furthermore, blood MS does not identify the origin and destination of secreted proteins. Also, MS proteomics of cell culture supernatants is not physiological (because it is performed *in vitro* and, frequently, in serum-free media (*7*)) and cannot identify destinations of factors. To overcome these limitations, we developed a method whereby all secreted proteins from a specific tissue are labeled by biotin, then collected by affinity-enrichment from distal organs and identified by quantitative MS (**Fig. 1A**). In this way, we simplify the proteome under study to the most relevant protein candidates involved in interorgan trafficking, and determine their origin and destination.

**Fig. 1:**
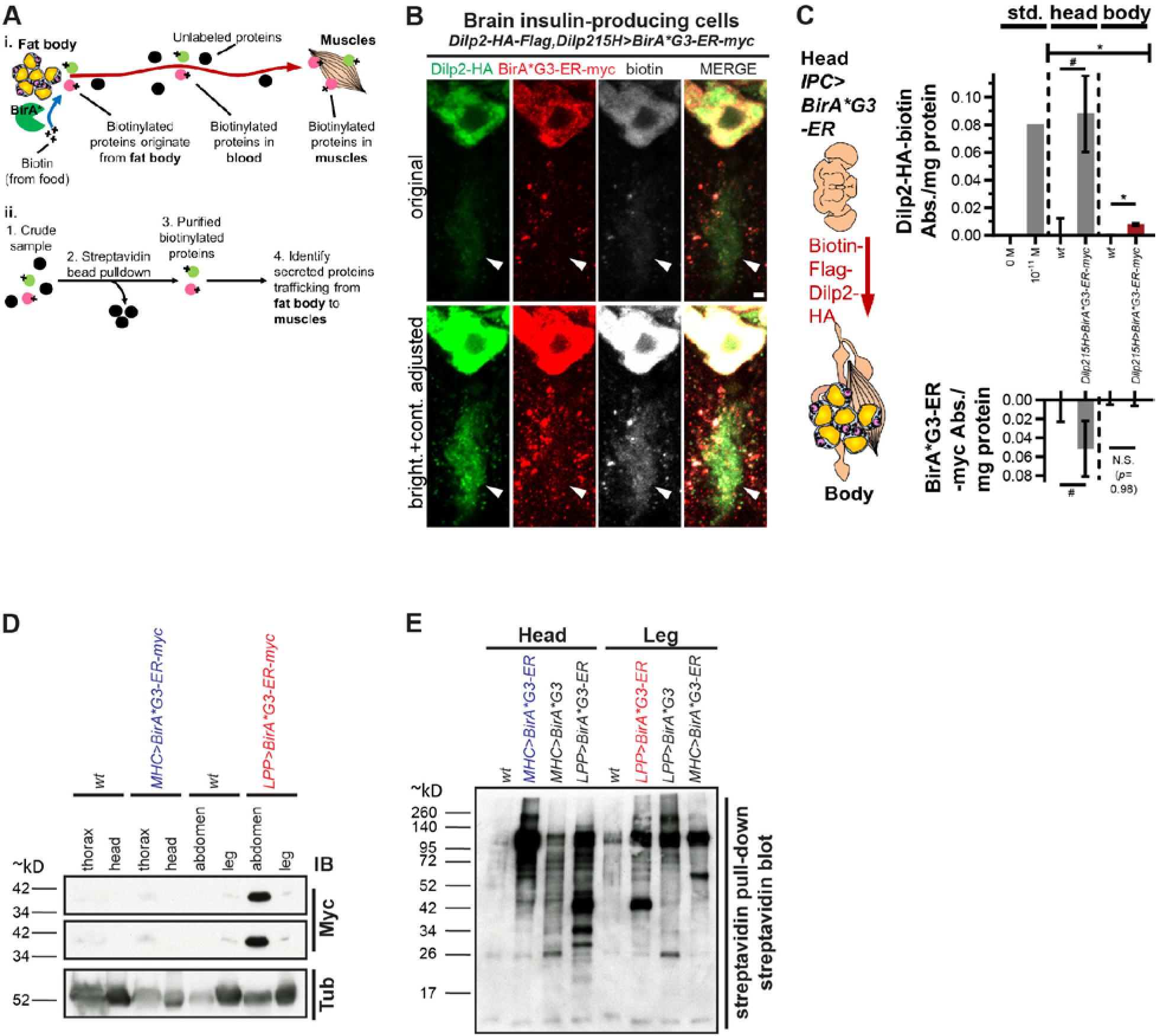
Labeling of secreted proteins and their detection in distal organs. **(A)** *i.* Promiscuous biotin ligase BirA*R118G or our newly-engineered high-activity BirA*G3 biotinylates all proteins in a subcellular compartment of one organ and biotinylated proteins traffic to other organs or body parts. *ii.* Biotinylated proteins are purified and identified by mass spectrometry. **(B)** *BirA*G3-ER* (endoplasmic reticulum) was expressed in insulin-producing cells (IPCs). BirA*G3-ER^+^, Biotin^+^, Dilp2-HA^+^ (*Drosophila* insulin-like peptide 2) (*12*) cell bodies, and BirA*G3-ER^negative^, biotin^+^ and Dilp2-HA^+^ projections (arrows) were detected. Scale bar: 2 μm. **(C)** Spatially informed bead enzyme-linked immunosorbent assay (ELISA) of biotinylated Dilp2-HA trafficking from head. Data is mean±SEM absorbance (Abs.). Top: *n*=2 biological replicates; one-way ANOVA (\**p*=0.037) and Holm-Sidak test (#*p*=0.10, \**p*=0.022). Std.: 1.04×10^−11^ M standard biotin-Flag-GS-HA peptide (*n*=1). Bottom: *n*=3 biological replicates; two-tailed t-test (#*p*=0.08996). N.S. means not significant. **(D)** Fat body (FB)-expressed *LPP-Gal4>BirA*G3-ER-myc* is detected in abdomens but not legs, and muscle-expressed *MHC-Gal4>BirA*G3-ER-myc* is detected in thoraces but not heads. Middle panel: higher exposure of the myc blot. IB: immunoblot. **(E)** Streptavidin bead pull-down followed by streptavidin-HRP detection in leg and head lysates after FB and muscle biotinylation using BirA*G3-ER or BirA*G3 (cytoplasmic/nuclear for unconventionally-secreted proteins) In **(B to E)** flies were maintained with 50 μM biotin in food during adulthood. *wt* means *wild-type*. See also: **Fig. S1 to S4**.

Biotinylation by the *E.coli* promiscuous biotin protein ligase (*BirA\**; “*” means promiscuous), *BirA*R118G* mutant (*8, 9*), or our newly-engineered high-activity *BirA*G3* (third generation evolved mutant, a relative of TurboID (*10*)), is continuous and mild, only requiring addition of biotin to culture media (*9*), potentially allowing long-term protein fate tracking. We developed the method in *Drosophila*, as organ systems and many physiological processes are conserved between flies and mammals. For instance, *Drosophila* has organs including the fat body (FB; functional equivalent to the mammalian liver and adipose tissue) and muscles, and systemic factors with mammalian orthologs, including insulin, leptin, glucagon, transforming growth-factor/activin-like, and tumor necrosis-factor (*1*). We generated *Drosophila* strains carrying *UAS-BirA*R118G-ER* and *UAS-BirA*G3-ER* to express BirA* in the endoplasmic reticulum (ER) for biotinylation of conventionally-secreted proteins. In addition, to detect unconventionally-secreted proteins (*2*), we generated untagged *UAS-BirA*R118G* and *UAS-BirA*G3* lines for biotinylation of the cytoplasm/nucleus. When expressed using a tissue-specific *Gal4* and biotin feeding, BirA* can cause biotinylation in the ER or cytoplasm/nucleus of any chosen organ. Expression of *UAS-BirA*G3-ER* using ubiquitous *TUB-Gal4* or FB-specific *LPP-Gal4* drivers (*11*) and biotin feeding did not have consistent, statistically-significant effects on lifespan or climbing ability (**Fig. S1**), suggesting that biotinylation does not severely affect organismal function, thereby enabling long-term studies.

As a proof-of-concept, we expressed *UAS-BirA*G3-ER* using the brain insulin-producing cell (IPC)-specific *Dilp215H-Gal4* (*12*) driver (Dilp2: *Drosophila* insulin-like peptide 2) and detected overlapping Dilp2-HA (*12*), BirA*G3-ER-myc, and biotinylated proteins in the brain (**Fig. S2, A to H**). Interestingly, we observed BirA*G3-ER-negative, Dilp2- and biotin-positive areas in IPC soma and projections, suggesting that Dilp2 traffics from BirA*G3-ER-positive production sites within cells (**Fig. S1B**). Further, using bead enzyme-linked immunosorbent assay (ELISA; **Fig. S2, I and J**) we detected biotinylated Dilp2-HA in heads and body, and BirA*G3-ER-myc in heads but not body (**Fig. 1C**). This suggests that the biotinylated protein (Dilp2-HA), but not the biotinylating enzyme (BirA*G3-ER), is secreted into the body. We conclude that our approach can sensitively detect low-abundance interorgan communication peptides such as Dilp2, which circulates on the order of 10^−10^ mol/L (*12*), and that biotinylation does not severely inhibit protein secretion.

To further verify that we can controllably express BirA*G3-ER in different tissues without undesirable secretion of the enzyme, we expressed *LPP-Gal4>BirA*G3-ER* in FB and detected BirA*G3-ER in abdominal FB (**Fig. S3**), but not legs (**Fig. 1D)**. In addition, we expressed *BirA*G3-ER* in muscles (*MHC-Gal4>BirA*G3-ER)* and detected BirA*G3-ER in thoraces (which contain muscles) but not heads (**Fig. 1D**). We observed that in legs, biotinylation patterns in *wild-type* (*wt*), *LPP-Gal4>BirA*G3-ER* (labels proteins secreted from ER), *LPP-Gal4>BirA*G3* (labels cytoplasmic/nuclear unconventionally-secreted proteins), and *MHC-Gal4>BirA*G3-ER* (leg muscles resident proteins) differed. Differences in biotinylation were also observed in heads, between *wt*, *MHC-Gal4>BirA*G3-ER*, *MHC-Gal4>BirA*G3*, and *LPP-Gal4>BirA*G3-ER* (head FB resident proteins) (**Fig. 1E**). Therefore, FB-secreted proteins found in legs, muscle-secreted proteins found in head, and ER-derived (*BirA*G3-ER*) and cytoplasm-derived (*BirA*G3*; non-classical/unconventional secretion) proteins differ significantly, consistent with organs’ (*1*) and organelles’ (ER versus cytoplasm) (*2*) secretome differences. Furthermore, using the original, less active *R118G* biotin ligase *BirA*R118G* and *BirA*R118G-ER*, we detected FB-secreted proteins in legs, muscle-secreted proteins in head, FB- and muscle-secreted proteins in hemolymph (fly blood), and FB-secreted proteins in brain, without detecting BirA*R118G or BirA*R118G-ER in distal tissues (**Fig. S4**). Altogether, these results demonstrate that the BirA* approach is able to detect secreted proteins in distal organs, and show its usefulness for identifying candidate proteins that traffic between multiple organs.

We used *BirA*G3-ER* and quantitative tandem mass-tag (TMT) MS to identify, in a single experiment, FB- and muscle-derived proteins in legs and heads, respectively (**Fig. S5**). We focused on this model organ system because: legs can be collected in large numbers (*11*), these interorgan communication axes are poorly-characterized, and FB-derived proteins present in legs may be involved in metabolic regulation of muscle activity (*1*). MS-identified proteins in legs were first compared to positive-control (PC) secreted/receptor and negative-control (NC) intracellular lists (**Fig. S6A**) (*11*). For each of the four *BirA*G3-ER/wild-type* (*wt*) TMT-ratio comparisons, we determined threshold TMT ratios for hit-calling at which 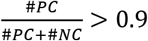 (**Fig. S6B**). From this, for each protein, we calculated an “MS-score”, defined as the number of TMT-ratio comparisons in which the protein TMT-ratio exceeded threshold (*11*). Hence, proteins with higher average TMT-ratios have higher MS-scores, highest-confidence hits have MS-scores of 4, background proteins (non-hits) have MS-scores of 0 (**Fig. 2A**; **Fig. S6C**), and as expected, hits are enriched for PC proteins (**Fig. S6D**). The biological replicates show good agreement in TMT-ratio signals (**Fig. 2A**).

**Fig. 2:**
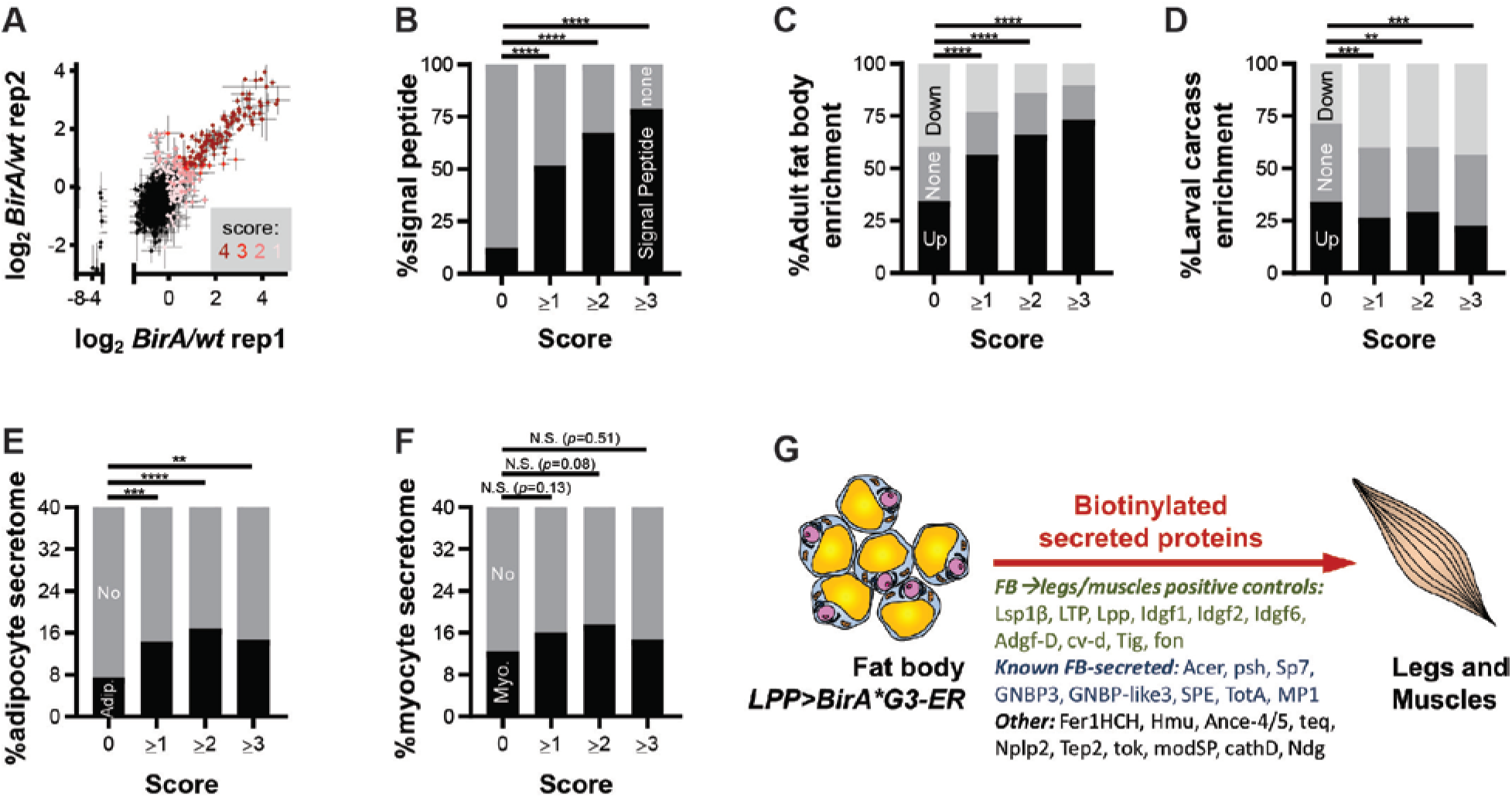
Identification of fat body (FB)-derived proteins in legs/muscle. FB was labeled using *LPP-Gal4>BirA*G3-ER* (endoplasmic reticulum) and biotinylated proteins from legs were analyzed by tandem mass-tag (TMT) mass spectrometry (MS). *Wt* (*wild-type*) legs were used as controls. Flies were maintained with 50 μM biotin in food during adulthood. **(A)** Leg log_2_(*BirA*G3-ER/wt*) TMT-ratios in two replicates. Each point is *n*=2 comparisons, mean±SEM log_2_TMT ratio. MS score: number of comparisons (from 4) in which TMT-ratio>threshold (score 4 is for most confident hits and 0 is background). **(B to F)** Statistics: chi-square test. N.S. means not significant. **(B)** As MS score increases, the fraction of proteins with putative signal peptides (*11*) increases. \*\*\*\**p*<0.0001. **(C and D)** As MS score increases, fraction of proteins enriched for adult FB mRNA microarray(*13*) expression increases **(C)**, and fraction of proteins enriched for larval carcass (FB-free muscle dataset) FB mRNA microarray(*13*) decreases **(D)**. \*\*\*\**p*<0.0001, \*\*\**p*=0.0003 (0 vs 1), \*\*\**p*=0.0008 (0 vs 3), \*\**p*=0.0055. **(E and F)** Hits are enriched for mammalian adipocyte **(E)** but not myocyte **(F)** secretome orthologs (*11*). \*\*\*\**p*<0.0001, \*\*\**p*=0.0004, \*\**p*=0.0090. **(G)** Model of identified proteins known to traffic from FB to imaginal discs or muscles, and known and novel FB-secreted proteins (*11*). See also: **Fig. S5 to S14**, **Tables S1 to S5**.

Our analysis revealed that higher MS-scores correlate significantly with more signal peptide-containing proteins, suggesting that our hits are secreted (**Fig. S6E**; **Fig. 2B**) (*11*). Consistently, hits are enriched for proteins found in whole fly hemolymph MS (**Fig. S7**; **Table S1**). Predicted transmembrane domain-containing proteins are also enriched in hits, consistent with shedding of known transmembrane factors (*1, 11*) (**Fig. S8A**). As negative controls, known ER-resident and predicted unconventionally-secreted proteins were not enriched (**Fig. S8, B to C**) (*11*). We estimated protein abundances using the PAX database (*11*), and determined that increased scores showed enrichment for lower whole-body abundance proteins (**Fig. S9**), likely reflecting the isolation of the most-relevant proteins away from abundant contaminants using biotinylation.

Interestingly, as MS-score increases, the fraction of hits that were enriched for adult FB mRNA microarray expression (*13*) increases (**Fig. 2C**). Consistently, overlaps with larval FB microarray and larval/pupal FB RNAseq datasets also increase (data not shown) (*11*). In addition, increased MS-scores result in decreased enrichment in larval carcass mRNA (FB-free muscle dataset (*13*); **Fig. 2D**). Thus, our dataset is enriched for genes transcribed in FB but not in muscles, suggesting that these proteins are candidates for trafficking from FB to legs/muscles. Moreover, hits are enriched for mammalian adipocyte secretome orthologs, but not for myocyte secretome orthologs (**Fig. 2, E and F**) (*11*), suggesting that some of our hits may be orthologs of previously-identified adipocyte secreted proteins.

Similar MS analysis identified muscle-secreted proteins harvested from heads (**Fig. S5 and S10**), and comparison with FB-to-legs dataset revealed limited overlap, suggesting organ specificity in interorgan secretome trafficking (**Fig. S11**). Furthermore, hits from the FB-to-legs *LPP-Gal4>BirA*G3-ER* dataset show statistically-significant overlap with hits from similarly-analyzed *LPP-Gal4>BirA*R118G-ER* MS dataset (first generation, less active BirA*; **Fig. S12**), compared to background (**Fig. S13**).

Overall, using *LPP-Gal4>BirA*G3-ER*, we identified 269 FB ER-derived proteins (corresponding to 266 genes) present in legs, including growth factors, binding-proteins, enzymes, peptidases, secreted kinases (**Fig. S14**), and 98 proteins of poorly-characterized functions (“Computed Genes”/CGs). We were able to detect FB-to-legs positive controls known to traffic from FB to muscles and/or imaginal discs, including lipid binding proteins Lsp1β, LTP, Lpp; growth-factors Idgf1/2/6 and Adgf-D; TGF-β binding protein cv-d; extracellular-matrix protein Tig; and the muscle adhesion protein fon (**Fig. 2G**; details and references in **Table S2**). For instance, the FB-secreted fon has been previously detected in muscles and shown to regulate their functions (*14*). Moreover, we identified additional secreted proteins previously unknown to be produced by FB or to traffic to legs or muscles (**Fig. 2G**; **Tables S3 and S4**). Hence, our data shows that we are able to identify known and novel FB-specific proteins present in legs. Notably, 60-70% of all hits and 50-60% of proteins with poorly-characterized functions (CGs) are well-conserved to humans (DIOPT score of at least 3 (*15*)). We identified *Drosophila* orthologs of human secreted proteins involved in systemic pathways, including ACE, APOB, TLL1, BMP1, LNPEP, TRHDE, ENPEP, ECE1, MME/NEP, KLKB1, CD109, and CTSD (details and references in **Table S5**). Characterization of novel orthologs will provide future insights into mammalian physiology.

Climbing-ability assays and immunohistochemical readouts of autophagy through extent of ubiquitin- and p62/ref(2)p-positive protein-aggregation are established indicators of *Drosophila* adult muscle function (*16*). Thus, we used FB-specific *LPP-Gal4>RNAi* to determine if candidates from our FB-to-legs MS list have effects on muscle function. We detected three proteins with putative signal peptides (*11*) in our FB-to-legs dataset – CG4332, CG2145, and CG31326 – whose knockdown in FB using two independent RNAi-lines each resulted in muscle phenotypes of poor climbing-ability and increased area of protein-aggregates (**Fig. 3**; **Fig. S15**; **Table S6**). Moreover, *CG31326*-FB-RNAi specifically in adults using *tub-gal80ts,LPP-Gal4* resulted in similar muscle-phenotypes (data not shown). As controls, we tested FB-phenotypes of FB-RNAi of *CG4332* and *CG2145* and did not find significant effects on FB lipid-droplets (**Fig. S16**). In addition, we used *DMEF2-Gal4* to perform RNAi of these factors in muscles, and did not observe consistent, statistically-significant climbing ability defects (**Fig. S17**), suggesting that for climbing activity, muscles are not a significant source of CG4332, CG2145, and CG31326. Interestingly, *CG2145* and *CG31326-mRNA* are enriched in adult-FB over muscles (larval-carcass) (*13*).

**Fig. 3:**
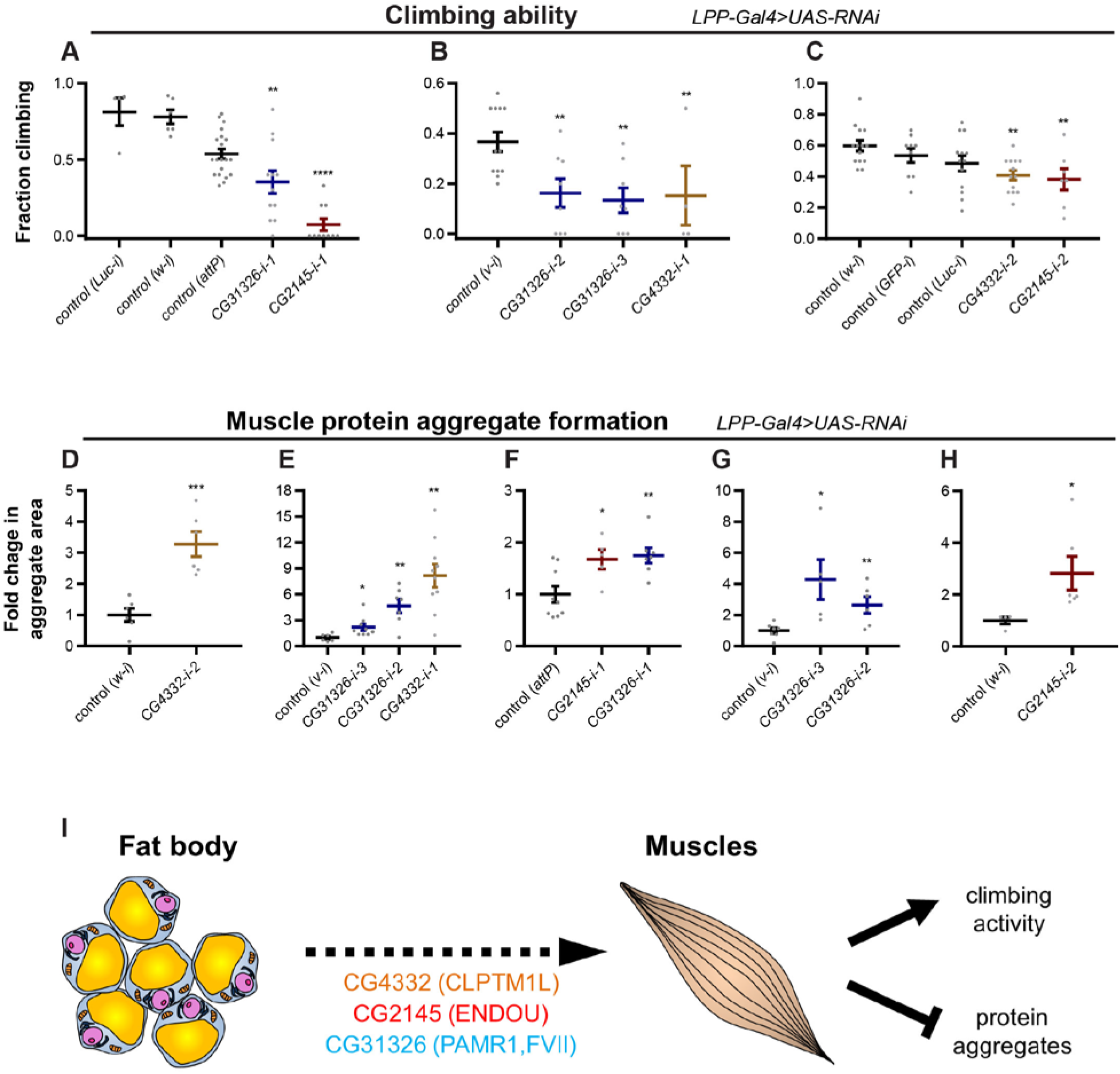
Muscle functions of legs/muscle proteins derived from fat body (FB). **(A to C)** Adult flies with FB RNAi (using *LPP-Gal4*) against *CG4332*, *CG2145*, and *CG31326* have reduced climbing ability. **(A)** *LPP-Gal4>RNAi* from VDRC (Vienna *Drosophila* Resource Center) at 3 weeks old and 29°C. Biological replicates: *n*=4 (*Luc-i*), *n*=6 (*w-i*), *n*=20 (*attP*), *n*=12 (*CG31326-i-1*), *n*=10 (*CG2145-i-1*). Relative to control (*attP*): \*\**p*=0.0024, \*\*\*\**p*<0.0001. **(B)** *LPP-Gal4>RNAi* from NIG (Japan National Institute of Genetics) at 5 weeks old and 27°C. Biological replicates: *n*=12 (*v-i*), *n*=8 (*CG31326-i-2*), *n*=8 (*CG31326-i-3*), *n*=4 (*CG4332-i-1*). Relative to control (*v-i*): \*\**p*=0.0037 (*CG31326-i-2*), \*\**p*=0.0011 (*CG31326-i-3*), \*\**p*=0.0090 (*CG4332-i-1*). **(C)** *LPP-Gal4>RNAi* from TRiP (Harvard Transgenic RNAi Project) at 5 weeks old and 27°C. Biological replicates: *n*=14 (*w-i*), *n*=9 (*GFP-i*), *n*=13 (*Luc-i*), *n*=13 (*CG4332-i-2*), *n*=7 (*CG2145-i-2*). Relative to control (*w-i*): \*\**p*=0.0021, \*\**p*=0.0031. Statistics: mean±SEM; one-way ANOVA and Benjamini, Krieger, Yekutieli Linear Two-Stage Step-Up FDR. **(D to H)** Adult flies with FB RNAi (using *LPP-Gal4*) against *CG4332*, *CG2145*, and *CG31326* have increased muscle protein aggregate formation. Area of p62/ref(2)p-positive protein aggregates was quantified, and normalized and compared to controls. **(D)** *LPP-Gal4>RNAi* (TRiP), 3 weeks old at 27°C. *n*=6 (*w-i*), *n*=6 (*CG4332-i-2*). \*\*\**p*=0.00052. **(E)** *LPP-Gal4>RNAi* (NIG), 3 weeks old at 27°C. *n*=6 (*v-i*), *n*=9 (*CG31326-i-3*), *n*=7 (*CG31326-i-2*), *n*=10 (*CG4332-i-1*). \**p*=0.029, \*\**p*=0.0017 (*CG31326-i-2*), \*\**p*=0.0011 (*CG4332-i-1*). **(F)** *LPP-Gal4>RNAi* (VDRC), 3 weeks old at 29°C. *n*=9 (*attp*), *n*=5 (*CG2145-i-1*), *n*=7 (*CG31326-i-1*). \**p*=0.021, \*\**p*=0.0046. **(G)** *LPP-Gal4>RNAi* (NIG), 5 weeks old at 27°C. *n*=7 (*v-i*), *n*=5 (*CG31326-i-3*), *n*=6 (*CG31326-i-2*). \**p*=0.012, \*\**p*=0.0094. **(H)** *LPP-Gal4>RNAi* (TRiP), 3 weeks old at 27°C. *n*=4 (*w-i*), *n*=6 (*CG2145-i-2*). \**p*=0.038. Statistics: mean±SEM; two-tailed t-test. **(I)** Our data suggests that we identified novel FB-derived factors that affect muscle function: CG4332 (human ortholog: CLPTM1L, cleft-lip and palate-associated transmembrane protein 1-like), CG2145 (human ortholog: ENDOU, poly-U-specific placental endonuclease), and CG31326 (human orthologs: PAMR1, peptidase domain-containing associated with muscle regeneration-1; and coagulation factor FVII). See also: **Fig. S15 to S17**, **Table S6**.

A putative human ortholog of CG4332, is CLPTM1L (cleft-lip and palate-associated transmembrane protein 1-like), which has no known functions in muscles or adipose tissue. DNA-sequence and expression-level variations of this gene have been associated with tumor division and apoptosis (*17–19*). Moreover, a putative human ortholog of CG2145 is ENDOU (poly-U-specific placental endonuclease), which is a signal-peptide-containing RNA-binding protein of uncharacterized function in muscles or adipose-tissue (*20–22*). Furthermore, putative human orthologs of CG31326 are PAMR1 (peptidase domain-containing associated with muscle regeneration-1) and coagulation factor FVII (**Table S6**). The function of PAMR1 is uncharacterized in muscles or adipose-tissue, but its expression is associated with muscle regeneration (*23*), and overexpression may decrease growth of breast cancer cells *in vitro* (*24*). Also, FVII is associated with fatty-diet feeding, obesity, insulin-resistance, and type-2 diabetes, and may be secreted by the liver and adipose-tissue (*25, 26*). Altogether, our results suggest that CG4332, CG2145, and CG31326 are candidate conserved FB-derived proteins that affect climbing ability and muscle protein aggregate formation (**Fig. 3I**; **Table S6**).

In conclusion, we have established a platform, using promiscuous BirA*, to investigate secreted protein trafficking from subcellular-compartments of specific cells to distal-organs *in vivo*. Through biotinylation of ER proteins in IPCs (Dilp2), FB, and muscle, we demonstrated labeling specificity and sensitivity, long-term monitoring, and wide-applicability of the method. Our platform provides biochemical evidence suggestive of proteins trafficking to other organs or distal body parts, and allows differentiation between ER- and non-conventionally-secreted proteins. Interestingly, our results indicate that the interorgan communication network of secreted proteins is more extensive than previously thought. We provide a resource for novel conserved secreted proteins that are candidate adipokines targeting legs, and candidate myokines targeting heads. We used our resource to determine muscle functions of novel FB-secreted proteins. Our results suggest that the FB secretes CG31326, CG2145, and CG4332, reduction in levels of which deteriorates climbing ability and increases protein aggregate formation in muscles. These factors’ respective human orthologs – PAMR1/FVII, ENDOU, and CLPTM1L – have no previously known roles in adipose tissue to muscle communication. Further characterization of our identified putative adipokines in *Drosophila* and mammalian models may yield important insights into systemic physiology.

In interpreting these results, one must exercise caution as the repertoire of identified proteins may include those present in the hemolymph, as opposed to those present specifically within muscles or heads. Further tests are required to determine if the FB-derived factors affect muscle function through a direct mechanism. However, while this may be due to detection limit differences, 60% of our FB-derived proteins identified in legs have not been observed in our hemolymph MS (see **Fig. S7C**). Also, we observed proteins known to traffic from FB to muscles or imaginal discs, and FB-secreted proteins in dissecting buffer-washed brains. Improvements to the platform include tissue dissection and washing to minimize hemolymph content; comparing leg/muscles proteomes to FB-derived hemolymph proteomes; combining BirA* labeling in FB with a different post-translational protein labeling approach in muscles to identify FB-derived proteins making contact with muscles; and increasing the activity and decreasing the stability of BirA* for more potent and tissue-restricted labeling. Our platform is widely applicable to research in interorgan, local (within a tissue), or intracellular (e.g., within neuronal projections) protein trafficking, unconventional secretion, protein trafficking in co-culture systems, and to mouse or other organisms *in vivo*, under healthy or diseased conditions. In addition, transiently-secreted proteins or those released due to perturbations (such as diet, stress, tumors) (*1*) may be studied by inducing BirA* biotinylation using biotin for a limited time. Another application is to enable spatially informed ELISA, as demonstrated using Dilp2 (see **Fig. 1C**), for proteins with only one available antibody – desirable for small or difficult antigens.

## Acknowledgments

I.A.D. acknowledges generous financial support from NSERC PGS-M and PGS-D, Brigham and Women’s Hospital Osher Pre-doctoral fellowship, and Harvard Medical School (HMS) Department of Cell Biology Innovative Grant Program (IGP). We thank Aldina Mesic and the Microscopy Resources on the North Quad (MicRoN) core at HMS for excellent technical assistance. We thank Seung Kim, Andreas Brech, TRiP at HMS, NIG, VDRC, Bloomington *Drosophila* Resource Center, and Developmental Studies Hybridoma Bank for stocks and antibodies. N.P. is an investigator of the Howard Hughes Medical Institute (HHMI). We acknowledge the National Institutes of Health (NIH) grants 5P01CA120964 (J.M.A.) and 5P30CA006516 (J.M.A.). This work was supported by a HCIA grant from HHMI.

## Author Contributions

I.A.D. conceived the method. I.A.D. and N.P. designed and supervised experiments. I.A.D., D.W., N.D.U., L.M., T.S., R.Z., A.T., and J.M.A. performed experiments. I.A.D., D.W., Y.H., N.D.U., L.M., T.S., J.M.A., S.A.C., and N.P. analyzed data. T.B. and A.Y.T. generated BirA*G3, which was cloned into *Drosophila* by J.A.B.. I.A.D. and N.P. wrote the manuscript with input from all authors.

