## Supplementary Information for "Proteomics of protein trafficking by *in vivo* tissue-specific labeling"

**This PDF file includes:**

Materials and Methods

Figs. S1 to S17

Tables S1 to S6

References

### Materials and Methods

#### *Drosophila* stocks

The following fly stocks were generated as part of this study. *BirA*R118G* was generated by mutating R118 to G in the pDisplay-BirA-ER (endoplasmic reticulum) (IgK signal peptide-HA-BirA-KDEL; Addgene 20856) (*1*) using QuikChange mutagenesis kit (Agilent). For the improved, highly-active *BirA*G3-ER*, a BiP signal peptide (for targeting to ER (*2*)) and KDEL were added using PCR primers. Constructs were cloned using Gateway methods (*3*) (Thermo-Fisher Scientific) into *pENTR/D-TOPO* (Thermo Scientific), and *pWalium10roe* vector was used as a final destination vector (*4*). The constructs were injected into *nanos-ΦC31-integrase* expressing embryos into *attP40* or *attP2* sites, as previously described (*4*). For *UAS-BirA*R118G-ER-HA(attP40)*, the complete genotype is *w;10xUAS-IgK-HA-BirA*R118G-KDEL,w$^+$(attP40,y$^+$)*. For *UAS-BirA*R118G-ER-HA(attP2)*, the complete genotype is *w;;10xUAS-IgK-HA-BirA*R118G-KDEL,w$^+$(attP2,y$^+$)*. For *UAS-BirA*R118G-HA(attP40)*, the complete genotype is *w;10xUAS-HA-BirA*R118G,w$^+$(attP40,y$^+$)*. For *UAS-BirA*R118G-HA(attP2)*, the complete genotype is *w;;10xUAS-HA-BirA*R118G,w$^+$(attP2,y$^+$)*. For *UAS-BirA*G3-ER-myc*, the complete genotype is *w;10xUAS-BiP-myc-BirA*G3-KDEL,w$^+$(attP40,y$^+$)*. For *UAS-BirA*G3-myc*, the complete genotype is *w;10xUAS-myc-BirA*G3,w$^+$(attP40,y$^+$)*.

The following additional stocks were used. Where indicated, *w[1118] wt* (*wild-type*) control was used. For hemolymph MS, *Oregon-R* (*Ore$^R$*) *wt* flies were used. In **Fig. 1, B and C** and **Fig. S2, A to H**, flies of the genotype *UAS-BirA*G3-ER-myc/(If or CyO);Dilp215H-Gal4,dilp2$^1$,Dilp2-HA-Flag(attP2)* were used. Flies of the genotype *w[1118];UAS-Dcr-2;Dilp215H,dilp2$^1$,Dilp2-HA-Flag(attP2)* were a gift of Seung Kim (*5*). These *Dilp2-HA-Flag* flies were previously used to measure changes in concentrations of systemic Dilp2 (*5*). Also, *LPP-Gal4* (lipophorin-Gal4) is a fat-body (FB)-specific driver (*6, 7*) (data not shown). *MHC-Gal4* and *DMEF2-Gal4* drivers were used for muscle expression (*8-10*). *TUB-Gal4* (*11*) and *10xUAS-IVS-mCD8-GFP(attP40)* (*12*) (Bloomington 32186) were also used (**Fig. S1**).

The following *UAS-RNAi* lines from Vienna *Drosophila* Resource Center (VDRC) (*13*), Japan National Institute of Genetics (NIG), and Harvard Transgenic RNAi Project (TRiP) (*4*): Control (*Luc-i*) (*Luceferase RNAi*; JF01355) (TRiP); Control (*w-i*) (*white RNAi*; HMS00017) (TRiP); Control (*attP*)

(*attP*[*VIE-260B*]; 60100) (VDRC); Control (*v-i*) (*vermillion RNAi*; 2155R-1) (NIG); Control (*GFP-i*) (*GFP RNAi*; HMS00773) (TRiP); *CG31326-i-1* (51890GD) (VDRC); *CG31326-i-2* (31326R-2) (NIG); *CG31326-i-3* (31326R-1) (NIG); *CG2145-i-1* (14874GD) (VDRC); *CG2145-i-2* (HMJ23623) (TRiP); *CG4332-i-1* (4332R-3) (NIG); and *CG4332-i-2* (HMS01341) (TRiP).

#### ***Drosophila* culture**

Flies were cultured using standard methods using Perrimon lab food (*14*). Preliminary experiments using *BirA*R118G-ER* showed that maximum BirA*R118G-ER expression and activity occurs at 30-31°C and with 2 copies of BirA*R118G-ER together with 2 copies of *MHC-Gal4* (*BirA*R118G-ER(attP40)/UAS-BirA*R118G-ER(attP40);MHC-Gal4/MHC-Gal4*), or 3 copies of *BirA*R118G-ER* and 1 copy of *LPP-Gal4* (*UAS-BirA*R118G-ER(attP40)/UAS-BirA*R118G-ER(attP40);LPP-Gal4/UAS-BirA*R118G-ER(attP2)*) (data not shown). For *BirA*R118G* or *BirA*R118G-ER*, crosses were performed at 22-25°C. After collection, adults were moved to 30-31°C for 4 days (with food changes every 2 days). Adults were subsequently transferred to 50 µM biotin food (see below for preparation) at 30-31°C for 6 days (with food changes every 2 days) until processing/collection. For *BirA*G3* or *BirA*G3-ER*, crosses were performed at 25°C. After eclosion and collection at 22°C, adults were moved to 29°C for 4 days (with food changes every 2 days). Adults were subsequently transferred to 50 µM biotin food (see below for preparation) at 29°C for 6 days (with food changes every 2 days) until processing/collection. For RNAi experiments, crosses were performed at 25°C. After eclosion and collection, adults were transferred to 27°C (TRiP and NIG stocks) or 29°C (VDRC stocks), and were regularly transferred to fresh food until experiments.

#### **Biotin-containing fly food preparation**

Biotin stock solution (around 18 mM) was prepared from solid biotin (Sigma B4639) in water, with final pH adjusted to around 7.2, and stored at -20°C. To prepare fly food containing 50 µM biotin, regular lab fly food was boiled three times until consistently liquid and the temperature was 100°C. The food was then weighed in an empty beaker, and the volume was calculated using the approximate food density of 1.04 g/mL. Next, the food was cooled down to around 60-65°C and a final concentration of 50 µM biotin was added. The mixture was then stirred well using a blender, and poured into vials or bottles (enough to cover the surface). Finally, the food was dried for several hours to overnight under cheesecloth, bottles or vials were plugged, and stored at 4°C until use.

This concentration (50 µM) of biotin was chosen based on published *in vitro* data with BirA*R118G(*15*), on *in vitro* experiments with *BirA*R118G* and *BirA*R118G-ER* (data not shown), and because *in vivo*, higher concentrations of biotin (500 µM) did not cause an increase in biotin labeling over 50 µM based on experiments with *LPP>BirA*R118G*, *DMEF2>BirA*R118G*, *MHC>BirA*R118G*, *ACT(actin)-Gal4>BirA*R118G*, or *CG(FB driver)-Gal4>BirA*R118G* (data not shown).

**Lifespan or survival analysis**

Lifespan analysis was performed according to established assays (*10*). Flies were raised on normal food without biotin, and after eclosion and collection, adults (males and females separately) were transferred at 29°C to normal food (no biotin) or to food containing 50 µM biotin for the remainder of their lives. Flies were transferred to fresh food regularly, and lethality was counted.

**Preparation of *Drosophila* body parts, tissues, and dissections**

Previously-described methods (*16-18*) were used to prepare *Drosophila* body parts and tissues, with modifications. Fly whole bodies were flash-frozen in liquid nitrogen by transferring flies to a 15 mL centrifuge tube using a funnel (no $CO_2$), and stored at -80°C until use. During head and leg preparation, flies were kept on dry ice for as much as possible, and body parts were prepared in small batches. A sieve assembly consisting of a #25 710 µm metal sieve on top, #45 355 µm metal sieve in the middle, and 3 mm chromatography paper on the bottom (Whatman 3030-917). Please note that for legs, preliminary experiments using *LPP-Gal4>UAS-GFP* suggested that leg femur, tibia, and tarsus (that is, below the coxa) (*19, 20*) do not have detectable FB or GFP expression (data not shown).

Flies were vortexed 4 times for 15 seconds each, with a rest on dry ice in between. Vortexed flies were then decanted onto the top of the sieve assembly (710 µm), shaken, and mixed with a brush. Bodies without heads were found on the top of the 710 µm sieve. Where appropriate, bodies were examined for absence of other body parts, transferred to a microcentrifuge tube, processed (see below), or flash frozen and stored at -80°C. Further, the 355 µm sieve was shaken and mixed with a brush. Heads were found on top of the 355 µm sieve. Where appropriate, heads were examined for absence of other body parts, transferred to a microcentrifuge tube, processed, or flash frozen and stored at -80°C. Finally, on the bottom chromatography paper, legs (consisting of those parts below the coxa) were separated from other debris, transferred to a microcentrifuge tube, processed, or flash frozen and stored at -80°C.

Brains, abdomens, and thoraxes were dissected in cold PBS (Invitrogen) as previously described(*10, 14, 21*), and processed further (see below). For brains in **Fig. S4D**, any FB that was loosely attached to the brain was carefully and completely removed. For western blots, thoraxes and abdomens were separated using razor blades. For immunofluorescence imaging, each thorax was cut into two to four fragments to expose the muscles, while abdomens were separated from thoraxes, cut on the ventral side to expose the FB and other internal organs, and internal organs were removed.

Hemolymph was collected according to previously-described methods (*22*), with modifications. Less than 20 flies were processed at one time. On the $CO_2$ pad, flies were punctured using a tungsten needle (Fine Science Tools). Based on the pattern of *LPP-Gal4>UAS-GFP* expression, for FB-derived proteins, we selected GFP/FB-negative regions of the dorsal thorax (middle to anterior scutum), just lateral to the midline (between the midline and the dorsocentral line defined by large dorsocentral bristles and medial end of the transverse suture) (*23*). Also, based on the pattern of *MHC-Gal4>UAS-GFP* expression, for muscle derived proteins, we selected GFP-negative regions of the dorsal abdomen (A4 yellow abdominal segment (*24*)), just lateral to the midline. Punctured flies were transferred to a 0.5 mL microcentrifuge tube that contained a hole (pierced with a 25 gage needle). The tube with flies was then transferred to a 1.5 mL low-protein binding microcentrifuge tube (Eppendorf), containing 15 µL of phosphate buffered saline (PBS). The two-tube assembly was then centrifuged for 5 min at 5,000 rpm at 4°C. Next, the 0.5 mL microcentrifuge tube was taken out, and the hemolymph in PBS in 1.5 mL tube was centrifuged for 5 min at 5,000 rpm at 4°C. Further, supernatant was collected, transferred to another 1.5 mL tube, and centrifuged at 14,000×g for 15 min at 4°C. Finally, supernatant was collected, transferred to another 1.5 mL tube, flash-frozen, and stored at -80°C. For hemolymph MS proteomics, a slightly modified hemolymph extraction protocol was employed: thorax-punctured flies were transferred in bulk

(>20 flies per tube) to pierced 0.5 mL microcentrifuge tube and centrifuged twice for 4 min at 4,000 rpm at 4°C. The next steps are as described above.

**Protein lysate preparation**

Protein lysates were prepared as previously described, with modifications (*21, 25*). The lysis buffer was RIPA buffer (made in the lab, 50 mM Tris, pH=8.0, 150 mM NaCl, 0.1%SDS, 0.5% sodium deoxycholate, 1% Triton X-100; or commercial (Pierce)) supplemented with 1 mM benzamidine hydrochloride (VWR), 4 µM pepstatin (VWR), 100 µM PMSF (Sigma-Aldrich), and one cOmplete ULTRA Mini EDTA-free protease inhibitor tablet (Roche). Tissues in lysis buffer were combined with zirconium oxide beads (NextAdvance) and homogenized several times for 4 minutes on setting 9 using the Bullet Blender (NextAdvance), with additions of extra lysis buffer where necessary, and brief centrifugations in between. Additional lysis buffer was added where necessary, and samples were left on ice for 30 min. Next, samples were centrifuged for 15 min at 16,000×*g* at 4°C and supernatants were transferred to low protein binding tubes (Eppendorf). Total protein concentrations were measured using the BCA protein assay kit (Pierce), according to manufacturer's instructions. Protein samples were diluted to equal concentrations in one experiment using lysis buffer. Protein lysates were flash frozen and stored at -80°C.

**Streptavidin beads pulldowns**

Streptavidin pulldowns were performed as previously described, with modifications (*25*). Streptavidin magnetic beads (Pierce 88817) were washed (using a magnetic stand for separation) twice in lysis buffer (see above) and resuspended in lysis buffer. For pulldown-western blot experiments in **Fig. 1E**, 64 µg of protein was combined with 32 µL of beads. In **Fig. S4A**, 140 µg of protein (1.42 µg/µL in lysis buffer) was combined with 40 µL of beads. In **Fig. S4D**, 36 µg of protein (0.25 µg/µL) was combined with 18 µL of beads. For MS experiments, 4.8 mg (*BirA*R118G-ER*, at 3.98 mg/mL) or 8.5 mg (*BirA*G3-ER*, at 12.2 mg/mL) of total protein per sample was combined with 0.6 mL of beads. Samples were incubated overnight at 4°C with end-over-end mixing. On the next morning, beads were washed twice in lysis buffer, once in 2 M urea in 10 mM Tris (pH=8.0), and twice in lysis buffer. Please see the western blotting section and quantitative mass spectrometry section for further bead processing.

**Western blotting**

Western blots were performed according to standard protocols, with modifications (*25, 26*). Equal amounts of total protein were mixed with 1× loading buffer (62.5 mM Tris pH=6.8, 2% sodium dodecyl sulfate, 10% glycerol, 0.025% bromophenol blue, 100 mM DTT; or ProSieve ProTrack Dual Color Protein Loading Buffer, Lonza) and boiled for 5 minutes. To elute bound biotinylated proteins in streptavidin pulldown experiments (see above), washed beads were resuspended in equal volumes of 1× loading buffer (diluted in lysis buffer), boiled for 5 min, and vortexed. Equal amounts of sample were loaded to a PAGEr EX gel (Lonza) and ran using ProSieve EX Running Buffer (Lonza) according to manufacturer's instructions. Spectra Multicolor Broad Range Protein Ladder (Pierce) was used as molecular weight marker. The samples on the gel were transferred to a nitrocellulose membrane using ProSieve EX transfer buffer (Lonza) for 20 min, according to manufacturer's instructions. The membrane was then rinsed 3 times in deionized water, followed by 3 times for 5 minutes each in TBST (Tris buffered saline (Sigma-Aldrich) with 0.1% Tween 20) with gentle shaking.

For streptavidin western blots, the membrane was incubated in blocking buffer (5% dialyzed bovine serum albumin (BSA), diluted in TBST) overnight at 4°C with gentle shaking. Next, the membrane was incubated for 1 hour at room temperature in 1:40,000 streptavidin-HRP (Invitrogen) in blocking buffer. Subsequently, the membrane was washed three times quickly in TBST, followed by 6 times, 5 minutes each in TBST with gentle shaking. Detection was performed using ECL hyperfilm (GE Healthcare), first using the ECL substrate (Pierce), then by thrice washing in deionized water and twice for 5 minutes each time in TBST, followed by the Supersignal West Pico substrate (Pierce).

For other western blots, the membrane was blocked in 5% milk in TBST for 1 hour at room temperature with gentle shaking. Next, the membrane was incubated in a primary antibody dilution in 5% milk in TBST overnight at 4°C with gentle shaking. Primary antibodies were: 1:1,000 mouse anti-myc (clone 9E10, Santa Cruz Biotechnology), 1:2,000 rat anti-HA (clone 3F10, Roche), 1:10,000 mouse anti-tubulin (clone B-5-1-2, Sigma-Aldrich; incubation can also be done for 1 hour at room temperature). Further, the membrane was washed three times quickly in TBST, followed by 6 times, 5 minutes each in TBST with gentle shaking. The membrane was then blocked in 3% milk in TBST for 1 hour at room temperature with gentle shaking, and incubated for 1-2 hours at room temperature in 3% milk in TBST with 1:5,000 to 1:10,000 dilution of HRP-conjugated secondary antibody: sheep anti-mouse (GE Healthcare), donkey anti-mouse (Jackson ImmunoResearch), or goat anti-rat (GE Healthcare). For tubulin western blot, a 1:10,000 secondary antibody dilution was used. Detection was performed using ECL hyperfilm (GE Healthcare), first using the ECL substrate (Pierce), followed by the Supersignal West Pico substrate (Pierce), and by Supersignal West Femto substrate (Pierce). Where necessary, the membrane was stripped using the Restore Plus Western Blot Stripping Buffer (Pierce) for 15 min at room temperature with gentle shaking, followed by thrice washing in deionized water, and 6 times, 5 minutes each washing in TBST.

**Fig. 1D** is a representative result of 3 western blots. Note that the thorax (*MHC-Gal4*) BirA*G3-ER-myc signal is lower than abdomen (*LPP-Gal4*) BirA*G3-ER-myc. However, thoracic BirA*G3-ER-myc is consistently detected in 3 western blots. **Fig. 1E** is a representative result of 2 western blots. In **Fig. 1, D and E** males and females were used, and *w[1118]* flies were used as *wt* control.

**Spatially-informed, one-antibody bead enzyme-linked immunosorbent assay (ELISA)**

By modifying standard ELISA methods (*5, 27*), we developed a spatially-informed, one-antibody bead ELISA protocol for biotinylated Dilp2-HA-Flag (schematic in **Fig. S2Q**). Protein lysates from heads and bodies from males and females were prepared as described above (**Preparation of *Drosophila* body parts, tissues, and dissections** and **Protein lysate preparation** sections). Standard Flag-GS-HA peptide (DYKDDDDKGGGGSYPYDVPDYA-$NH_2$, molecular weight 2411.45 Da, LifeTein) (*5*) was dissolved in water, aliquoted, and stored at -80°C. Note that the standard peptide concentration calculation took into account synthesized peptide purity. Flag-GS-HA peptide biotinylation was performed with EZ-link sufo-NHS-biotin (Thermo Scientific), according to manufacturer’s instructions. Flag-GS-HA (final concentration of 2.5 mg/mL) or water background control was combined with 5-, 20-, or 50-fold molar excess of sulfo-NHS-biotin in a total reaction volume of 6 µL for at least 2 hours on ice. For the standard peptide in **Fig. 1C**, we used a 20-fold excess of sulfo-NHS-biotin. Biotinylated Flag-GS-HA was stored at -80°C or used. Biotinylated Flag-GS-HA or background control dilutions for ELISA were performed in fetal bovine serum (FBS, heat-inactivated, Gibco).

We coupled antibodies to tosylactivated Dynabeads M-280 (Invitrogen), based on manufacturer's instructions. Beads were washed twice 500 µL of 0.1 M borate buffer, pH=9.5 (using a magnetic stand for separation) in a 0.5 mL low-protein binding tube (Eppendorf). Next, 1 mg of beads, 20 µL of borate buffer, 40 µL of 3 M ammonium sulfate in borate buffer, and 40 µL of 0.5 mg/mL of antibody were combined (these amounts were scaled as needed), and incubated end-over-end at 37°C overnight. The antibodies used were: rat anti-HA (clone 3F10, Roche) and mouse anti-myc (clone 9E10, Santa Cruz Biotechnology). For the anti-myc antibody, the original concentration was 0.2 mg/mL, and it was concentrated to approximately 0.5 mg/mL using a 30 kD cutoff centrifugation concentration device (Amicon, EMD Millipore). Alternatively, 1 mg of beads, 65.88 µL of 3 M ammonium sulfate in borate buffer, and 100 µL of 0.2 mg/mL antibody solution were combined, and incubated end-over-end at 37°C overnight. For biotinylated Dilp2-HA-Flag detection, we chose to not use anti-flag as the capture antibody because the flag tag (DYKDDDDK) (*28*) has a number of lysine groups which may be modified by biotin, potentially resulting in a loss of immunoreactivity.

Next, the beads were washed once with 1.5 mL of wash buffer (0.05% Tween-20 in PBS) and thrice with 1 mL of PBS. Then, beads were incubated with 1 mL of The Blocking Solution (Candor) for 1 hour at room temperature with end-over-end mixing. Subsequently, beads were washed twice with 1 mL of wash buffer. Next, beads were blocked in 1 mL of 3 M ethanolamine for at least 2 hours at room temperature, with end-over-end mixing. Beads were then washed thrice with 1 mL of wash buffer. Afterwards, the beads were split into individual low protein bind vials (Eppendorf). In each vial, 50 µg (**Fig. S2R**) or 100 µg (**Fig. 1C**) of blocked beads were combined with 1 mL of biotinylated Flag-GS-HA standard or tissue protein lysates (input total protein concentration: 1-4 mg/mL for head and 2-17 mg/mL for body in different experiments), and incubated overnight at 4°C with end-over-end mixing. In **Fig. S2R**, the background water control was diluted to the same extent as the highest concentration dilution of the Flag-GS-HA standard ($1.25\times10^{-9}$ M). Next, samples were washed thrice with 1 mL of wash buffer, and incubated with 100 µL of 1.25 µg/mL avidin-HRP (Affymetrix) in 1× ELISA/ELISPOT diluent (5× stock solution, Affymetrix) for around 30 min (**Fig. S2R**) to 1 hour (**Fig. 1C**) at room temperature with shaking. Samples were then washed at least 4 times in 1 mL of wash buffer, followed by once in 200 µL of wash buffer. Finally, samples were incubated with 100 µL 1× TMB substrate solution (Affymetrix) at room temperature with shaking for 12 minutes (HA ELISA, **Fig. 1C**), 15 min (myc ELISA, **Fig. 1C**), or 30 min (**Fig. S2R**). The reaction was terminated by adding 50 µL of stop solution (1 M $H_3PO_4$).

Sample absorbance values at $\lambda$=450 nm and $\lambda$=570 nm were measured using a spectrophotometer. In **Fig. 1C**, Nanodrop 8000 (Thermo-Fisher) was used, and in **Fig. S2R**, Spectramax Paradigm plate reader (Molecular Devices) was used. For biotinylated Flag-GS-HA standard, final absorbance (Abs.) values were calculated as: Abs.=$\lambda_{450\text{nm}} - \lambda_{570\text{nm}}$. For tissue samples, final Abs. values per mg of input protein were calculated as: Abs./mg protein=$\frac{\lambda_{450\text{nm}}-\lambda_{570\text{nm}}}{\text{mg input protein}}$. In **Fig. 1C**, normalization was performed by subtracting Abs. of 0 M standard peptide from Abs. of $1.04\times10^{-11}$ M standard peptide, average *wt* head Abs./mg protein from head Abs./mg protein, and average *wt* body Abs./mg protein from body Abs./mg protein.

Shown in top panel of **Fig. 1C** (Dilp2-HA-biotin ELISA) are representative results from two independent experiments. Shown in bottom panel of **Fig. 1C** (BirA*G3-ER-myc-biotin ELISA) are measurements from another similar head and body preparation of the same genotypes of flies. The three replicates from the same head and body preparation were measured across 2 runs. In **Fig. 1C**, *w[1118]* flies are used as *wt* controls.

**Quantitative tandem mass-tag (TMT) MS**

Female and male adult flies were used in BirA*R118G-ER and BirA*G3-ER MS experiments, and *w[1118]* flies were used as *wt* controls. Protein lysates and pulldowns were performed as described above (**Preparation of *Drosophila* body parts, tissues, and dissections**, **Protein lysate preparation**, and **Streptavidin beads pulldowns** sections). The different MS samples and their associated TMT labels are presented in **Fig. S5 and S12A**.

***On-bead digestion.*** Samples collected from *Drosophila melanogaster* legs and heads and enriched with magnetic streptavidin beads were washed twice with 200 µL of 50mM Tris-HCl buffer (pH 7.5), transferred into new 1.5 mL Eppendorf tubes, and washed 2 more times with 2 M urea/50 mM Tris (pH 7.5) buffer. Samples were incubated in 0.4 µg trypsin in 80 µL of 2 M urea/50mM Tris buffer with 1 mM DTT, for 1 hour at room temperature while shaking at 1000×*g*. Following pre-digestion, 80 µL of each supernatant was transferred into new tubes. Beads were washed twice with 60 µL of 2M urea/50 mM Tris buffer, and these washes were combined with the supernatant. The eluates were spun down at 5000×*g* for 1 min and the supernatant was transferred to a new tube. Samples were reduced with 4 mM DTT for 30 min at room temperature, with shaking. Following reduction, samples were alkylated with 10 mM iodoacetamide for 45 min in the dark at room temperature. An additional 0.5 µg of trypsin was added and samples were digested overnight at room temperature, while shaking. Following overnight digestion, samples were acidified (pH <3) with neat formic acid (FA), to a final concentration of 1% FA. Samples were spun down and de-salted on C18 StageTips as previously described(*29*). Eluted peptides were dried to completion and stored at -80°C.

***TMT labeling of peptides.*** Desalted peptides were labeled with TMT (10-plex) reagents (Thermo Fisher Scientific). Each TMT reagent was resuspended in 41 µL of MeCN. Peptides were resuspended in 100 µL of 50 mM HEPES and labeled with the TMT reagents as described in **Fig. S5 and S12A**. Samples were incubated at RT for 1 hr with end-over-end rotation. TMT reaction was quenched with 8 µL of 5% hydroxylamine at room temperature for 15 min with shaking. TMT labeled samples were combined, dried to completion, reconstituted in 100 µL of 0.1% FA, and desalted on StageTips.

***SCX stage tip fractionation of peptides.*** To increase depth-of-coverage, 50% of the TMT labeled peptide sample was fractionated by strong cation exchange (SCX) StageTips packed with 3 disks of SCX (3M Empore) material on top of 2 disks of C18 material. StageTips were conditioned with 100 µL of 100% MeOH, followed by 100 µL of 0.5% acetic acid/80% MeCN. Next, StageTips were equilibrated with 100 µL of 0.5% acetic acid, followed by 100 µL of 0.5% Acetic Acid/ 500mM NH4AcO/ 20% MeCN, and 100 µL of 0.5% acetic acid. Peptide samples were resuspended in 250 µL of 0.5% acetic acid and loaded onto the StageTips, washed 2x with 100 µL of 0.5% acetic acid, and transeluted from SCX material onto the C18 material with 100 µL of 0.5% acetic acid/80% MeCN. Three step-wise elutions from the C18 disks were completed as follows: the first fraction was eluted with 50 µL of 50 mM $NH_4AcO$; 20% MeCN (pH 5.15, adjusted with acetic acid), the second with 50 µL of 50 mM $NH_4AcO$: 20% MeCN (pH 8.25, adjusted with acetic acid), the third with 50 µL 50 mM $NH_4AcO$: 20% MeCN (pH 10.3, adjusted with acetic acid). Each eluate was collected separately and 200 µL of 0.5% acetic acid was added to each. Each fraction was desalted on C18 StageTips as described above, and samples were dried to completion.

***Liquid chromatography and mass spectrometry.*** Desalted peptides were re-suspended in 9 µL of 3% MeCN/0.1% FA and analyzed by online nanoflow liquid chromatography tandem mass spectrometry (LC-MS/MS) using a Q Exactive Plus mass spectrometer (Thermo Fisher Scientific) coupled on-line to a Proxeon Easy-nLC 1200 (Thermo Fisher Scientific) as previously described(*29*). Briefly, 4 µL of each sample was loaded at onto a microcapillary column (360 µm outer diameter × 75 µm inner diameter) containing an integrated electrospray emitter tip (10 µm), packed to approximately 24 cm with ReproSil-Pur C18-AQ 1.9 µm beads (Dr. Maisch GmbH) and heated to 50C. SCX fractionated samples were analyzed using a 110 min LC-MS method. Mobile phase flow rate was 200 nL/min, comprised of 3% acetonitrile/0.1% formic acid (Solvent A) and 90% acetonitrile /0.1% formic acid (Solvent B). The 110-minute LC-MS/MS method used the following gradient profile: (min:%B) 0:2; 1:6; 85:30; 94:60; 95:90; 100:90; 101:50; 110:50 (the last two steps at 500 nL/min flow rate). Non-fractionated samples were were analyzed using a 260 min LC-MS/MS method with the following gradient profile: (min:%B) 0:2; 1:6; 235:30; 244:60; 245:90; 250:90; 251:50; 260:50 (the last two steps at 500 nL/min flow rate). The Q Exactive was operated in the data-dependent mode acquiring HCD MS/MS scans ($r$ = 17,500) after each MS1 scan ($r$ = 70,000) on the top 12 most abundant ions using an MS1 target of $3 \times 10^6$ and an MS2 target of $5 \times 10^4$. The maximum ion time utilized for MS/MS scans was 120 ms; the HCD-normalized collision energy was set to 28; the dynamic exclusion time was set to 20 s, and the peptide match and isotope exclusion functions were enabled. Charge exclusion was enabled for charge states that were unassigned, 1 and >7.

**BirA* MS data analysis**

All protein trafficking MS data were analyzed using Spectrum Mill software package v 6.1 pre-release (Agilent Technologies). Similar MS/MS spectra acquired on the same precursor m/z within +/- 60 s were merged. MS/MS spectra were excluded from searching if they were not within the precursor MH+ range of 750-4000 Da or if they failed the quality filter by not having a sequence tag length >0. MS/MS spectra were searched against all *Drosophila melanogaster* proteins annotated at UniProt database (*30*) containing 21,979 proteins, and 259 common contaminants. All spectra were allowed +/- 20 ppm mass tolerance for precursor and product ions, 30% minimum matched peak intensity, and “trypsin allow P” enzyme specificity with up to 2 missed cleavages. The fixed modifications were carbamidomethylation at cysteine, and TMT at N-termini and internal lysine residues. Variable modifications included oxidized methionine and N-terminal protein acetylation. Individual spectra were automatically designated as confidently assigned using the Spectrum Mill autovalidation module. Specifically, a target-decoy based false-discovery rate (FDR) scoring threshold criteria via a two-step auto threshold strategy at the spectral and protein levels was used. First, peptide mode was set to allow automatic variable range precursor mass filtering with score thresholds optimized to yield a spectral level FDR of <1.2%. A protein polishing autovalidation was applied to further filter the peptide spectrum matches using a target protein-level FDR threshold of 0. Following autovalidation a protein-protein comparison table was generated, which contained experimental over control TMT ratios. For all experiments, non-*Drosophila* contaminants and reverse hits were removed. Furthermore, data was was median normalized.

Next, we established threshold TMT ratios for hit-calling using positive control (PC) secreted/receptor and negative control (NC) intracellular lists. For the PC list, we used fly receptor and secretome lists (*31*), as well as fly orthologs (using DIOPT (*32*)) of human receptome (*33*), human secreted proteins annotated at UniProt (*30*) and blood plasma MS (highly confident cumulative multiple-dataset PeptideAtlas list (*34*), and a list from humans with trauma (*35*)). PC proteins were also checked for presence of a signal peptide (*36*) and a transmembrane (TM) domains (*37*). For the NC list, we used high-confidence mitochondrial proteins (*26*), as well as cytoskeletal and high confidence transcription factors (nuclear) (*31*). Proteins identified by our *BirA*R118G-ER* and *BirA*G3-ER* MS were compared to the PC and NC lists, and assigned to as being a PC or NC protein (**Fig. S6A, S10B, S12B, and S12E**). For each experiment (head or leg), there were four TMT ratio comparisons (*BirA**-rep1/*wt*-rep1, *BirA**-rep1/*wt*-rep2, *BirA**-rep2/*wt*-rep1, *BirA**-rep2/*wt*-rep2; rep means replicate) (**Fig. S5 and S12A**). For each *BirA*G3-ER* comparison, we plotted the fraction $\frac{\#PC}{\#PC+\#NC}$ above each $\log_2$*BirA*/wt* TMT ratio, and determined a threshold TMT ratio at which $\frac{\#PC}{\#PC+\#NC}$ was generally greater than 0.9 (**Fig. S6B** and data not shown). Based on the shapes of the curves, the thresholds were: leg 127N/126 TMT-ratio≥0.43 at $\frac{\#PC}{\#PC+\#NC} > 0.93$; leg 127N/126 TMT-ratio≥0.5 at $\frac{\#PC}{\#PC+\#NC} > 0.92$; leg 129N/126 TMT-ratio≥0.7 at $\frac{\#PC}{\#PC+\#NC} > 0.95$; leg 129N/128C TMT-ratio≥0.7 at $\frac{\#PC}{\#PC+\#NC} > 0.93$; head 128N/127C TMT-ratio≥0.8 at $\frac{\#PC}{\#PC+\#NC} >$

$0.75$ to $0.8$ (the $\frac{\#PC}{\#PC+\#NC}$ vs 128N/127C TMT-ratio curve had significant fluctuation around the cutoff, and a manually-drawn best-fit curve had a $\frac{\#PC}{\#PC+\#NC}$ of at least around 0.8 to 0.9); head 128N/129C TMT-ratio≥0.9 at $\frac{\#PC}{\#PC+\#NC} > 0.91$; head 130N/127C TMT-ratio≥0 at $\frac{\#PC}{\#PC+\#NC} > 1.0$; and head 130N/129C TMT-ratio≥0.1 at $\frac{\#PC}{\#PC+\#NC} > 0.83$ (approximate; midpoint between TMT-ratio=0 at $\frac{\#PC}{\#PC+\#NC} = 0.67$ and TMT-ratio=0.2 at $\frac{\#PC}{\#PC+\#NC} = 1.0$). For *BirA*R118G-ER* the threshold TMT-ratios were set based on the shapes of the curves: leg 129C/127C TMT-ratio≥2.4 at $\frac{\#PC}{\#PC+\#NC} = 0.67$ (the next point is $\frac{\#PC}{\#PC+\#NC} = 1.0$); leg 129C/129N TMT-ratio≥1.4 at $\frac{\#PC}{\#PC+\#NC} = 0.86$; leg 127N/127C TMT-ratio≥3.4 at $\frac{\#PC}{\#PC+\#NC} = 1.0$; leg 127N/129N TMT-ratio≥2.8 at $\frac{\#PC}{\#PC+\#NC} = 1.0$; head 130C/128N TMT-ratio≥1.2 at $\frac{\#PC}{\#PC+\#NC} = 1.0$; head 130C/130N TMT-ratio≥0.8 at $\frac{\#PC}{\#PC+\#NC} = 1.0$; head 128C/128N TMT-ratio≥0.4 at $\frac{\#PC}{\#PC+\#NC} = 0.7$; and head 128C/130N TMT-ratio≥0.2 at $\frac{\#PC}{\#PC+\#NC} = 0.89$. MS-score was defined as the number of comparisons in which a specific protein's TMT ratio exceeds the threshold TMT ratio (**Fig. 2A; Fig. S10A, S12D, and S12G**). Thus, proteins with score of 0 are background, proteins with a score of 1 are lower-confidence hits, and proteins with a score of 4 are highest confidence hits. Note that in **Fig. 2A and Supplementary Figs. S6, A to C, E, S10, A, B, and D, S11A, and S12**, means (and where applicable, SEMs) were calculated from $\log_2$ values of TMT-ratios.

To obtain human ortholog information, we used DIOPT, versions 5 and 6 (*32*). MS-identified proteins at different scores were examined for the presence of putative signal peptides, using SignalP program (*36*) (**Fig. 2b; Fig. S6E, S10D, S10E**). Moreover, proteins were examined for their presence in our fly total hemolymph protein MS (**Fig. S7; Table S1**). Also, using TMHMM program(*37*), proteins were examined for the presence of transmembrane domains (**Fig. S8A and S10G**). In addition, based on UniProt annotation (*30*), we determined whether the identified proteins could be ER resident (**Fig. S8B and S10F**). Moreover, using SecretomeP program (*38*), we examined whether some of the proteins could be unconventionally/non-classically secreted (**Fig. S8C and S10H**). We also obtained tissue mRNA expression information from FlyAtlas microarray(*39*) (**Fig. 2, C and D; Fig. S10, I and J**) and RNAseq (*40*) (data not shown) datasets. In addition, protein abundance information was from integrated entire organism PAX database for *Drosophila melanogaster* (*41*) (**Fig. S9**). In **Fig. S14**, categorization of hits (score ≥ 1) into broad categories was performed manually using data from FlyBase (*42*) and NCBI Gene (*43*).

Further, we compared the MS-identified proteins to fly orthologs (using DIOPT version 5.3 (*32*)) of mammalian adipocyte (*44-53*) and myocyte (*54-61*) secretomes. For higher confidence, we required a protein to be identified in at least 2 adipocyte or myocyte secretome datasets to be considered an adipocyte or myocyte secreted protein for our analysis in **Fig. 2, E and F**.

**Hemolymph MS proteomics**

We performed whole hemolymph MS proteomics as part of our study to gain insight into which proteins may be circulating in *Drosophila* (**Tables S2 and S3**; **Fig. S7**). We identified a total of 1561 proteins, including 688 proteins of poorly-characterized functions ("Computed Genes"/CGs; FlyBase(*42*) r551). The following hemolymph MS experiments were performed: *Ore*$^R$ unfractionated hemolymph, *myoglianin* overexpressing (*21*) unfractionated hemolymph, methanol-extracted peptides from *Ore*$^R$ hemolymph, size-fractionated *Ore*$^R$ hemolymph, 4 week old *Ore*$^R$ fly unfractionated hemolymph, glycopeptide pull-down *Ore*$^R$ hemolymph, and non-tryptic (open) *Drosophila* database search of <3 kD fraction and O-glycopeptide pulldown experiment. In **Table S1** and **Fig. S7**, the MS data was analyzed by counting the number of times a peptide or different peptides for one protein in each of the above 7 experiments (for the purposes of this analysis, each of the above was counted as an experiment) were observed. Each identified protein was assigned to a PC secreted/receptor or NC intracellular lists as above (see **BirA* MS Data Analysis** section). Please note that for this analysis, if a protein was assigned to both NC and PC list, it was assigned to the "other" category. In addition, a targeted analysis was performed on *Ore*$^R$ unfractionated hemolymph for predicted or known (*62*) tryptic peptides; this analysis was not included in the above results.

We used methanol to precipitate proteins and enrich for smaller peptides (*63-65*). A total of 38 µL of hemolymph was precipitated in 80% methanol at -80°C for 7 hours. Next, the sample was centrifuged at 14,000×*g* for 15 min at 4°C, and supernatant was dried in a speed-vac for 2 hours.

Previously-established methods (*66*) were used for hemolymph size-fractionation. A total of 100 µL of hemolymph was dissolved in urea buffer (8 M urea, 50 mM Tris (pH=7.5), 75 mM NaCl) with 0.5%

dithiothreitol (DTT), and incubated at 60°C for 1 hour. Next, samples were incubated in 15 mM iodoacetamide for 30 min in the dark at room temperature with shaking, followed by 5 mM DTT for 15 min at room temperature in the dark. Next, we used Amicon Ultra 0.5 mL columns (Millipore) according to manufacturer's instructions. Columns (3K cutoff; Millipore) were pre-spun in urea buffer for 25 min at room temperature at 14,000×*g*. Next, the hemolymph sample was applied and centrifuged for 40 min at 14,000×*g* at room temperature. Further, 400 µL of urea buffer was applied and centrifugation was repeated. This was repeated twice more to obtain the <3 kD sample in the flow through. Next, the column was inverted into a new vial and centrifuged at 1,000×*g* for 2 min at room temperature. This was repeated four times with 100 µL of urea buffer to wash. The latter >3 kD fraction was transferred to pre-spun (with urea buffer at 30 min for 14,000×*g*) 10K membranes (Millipore), and centrifuged at 14,000×*g* for 30 min at room temperature. The columns were washed three times with 420 µL of urea buffer to obtain the 3-10 kD fraction. Further, to obtain the >10 kD fraction, the columns were inverted and eluted to new tubes by centrifuging at 1,000×*g* for 2 min at room temperature, followed by four times washing with 100 µL of urea buffer and centrifuging. The latter >10 kD fraction was transferred to pre-spun (with urea buffer at 18 min for 14,000×*g*) 30K membranes (Millipore), and centrifuged at 14,000×*g* for 15 min at room temperature. The columns were washed three times with 420 µL of urea buffer to obtain the 10-30 kD fraction. Further, to obtain the >30 kD fraction, the columns were inverted and eluted to new tubes by centrifuging at 1,000×*g* for 2 min at room temperature, followed by twice washing with 50 µL of urea buffer and centrifuging. The >30 kD sample was diluted 1:10 in 100 mM ammonium bicarbonate, pH=8.3 (final urea concentration was 0.8 M), 1 mM calcium chloride, and digested with sequencing grade modified trypsin (Promega) at 37°C overnight with shaking, according to manufacturer's instructions. Samples were then acidified with trifluoroacetic acid (TFA) to a final pH<2. The samples were subsequently purified using tC18 sep-pak columns (Waters) according to standard protocols (*67*). We used 200 mg cartridges for >10 kD fractions and 100 mg cartridges for <10 kD fractions. Columns were conditioned using 3 mL (200 mg cartridges) or 2 mL (100 mg) acetonitrile, followed by twice with 3 mL (200 mg) or 2 mL (100 mg) 50% acetonitrile in water and 0.5% acetic acid in water. Next, 3 mL (200 mg) or 2 mL (100 mg) of 0.1% TFA in water was passed through the column twice, and samples were slowly loaded. Columns were next washed twice with 3 mL (200 mg) or 2 mL (100 mg) of 0.1% TFA in water, followed by 360 µL of 0.5%

acetic acid in water. The samples were twice eluted with 0.9 mL of 50% acetonitrile with 0.5% acetic acid in water. Finally, the samples were dried in a speed-vac and frozen at -80°C.

Glycosylated peptide pull-downs were performed as previously described (*68-71*). Hemolymph (125 µL) was dissolved in a 0.4 M $NH_4HCO_3$ (pH=8.3), 0.1% sodium dodecyl sulfate (SDS), and 8 M urea. Next, 10 mM TCEP (Tris (2-carboxyethyl) phosphine)) was added and the mixture was incubated at 58°C for 1 hour 40 min. Then, 14 mM iodoacetamide was added, and incubated at room temperature for 30 min in the dark with mixing. This was followed by the addition of 5 mM TCEP for 15 min at room temperature in the dark. The sample was then diluted 1:10 in a final concentration of 100 mM $NH_4HCO_3$, pH=8.3 (final urea concentration was 0.8 M), 1 mM calcium chloride, and digested with sequencing grade modified trypsin (Promega) at 37°C overnight with shaking, according to manufacturer's instructions. The sample were then centrifuged for 12,000×*g* for 10 min and the supernatant were acidified with trifluoroacetic acid (TFA) to a final pH<2. The sample was subsequently purified using a tC18 sep-pak column (Waters). The column was conditioned using 3 mL of acetonitrile, followed by twice with 3 mL of 50% acetonitrile with 0.1% TFA in water. Next, 3 mL of 0.1% TFA in water was passed through the column twice, and the sample were slowly loaded. The column was next washed thrice with 3 mL of 0.1% TFA in water. The sample was twice eluted with 0.9 mL of 50% acetonitrile with 0.1% TFA in water. Subsequently, 10 mM sodium periodate was added and the sample was incubated for 1 hour in 4°C in the dark. Next, the 2 mL sample was diluted with 16.2 mL of 0.1% TFA in water and purified with a 200 mg tC18 sep-pak column (Waters) in the same way as described above, with the modification that the sample elution was with 0.1% TFA in 80% acetonitrile in water. Next, 300 µL of Affi-Gel hydrazide slurry (Biorad) was centrifuged at 3,000 rpm for 30 seconds, washed once with water, centrifuged at 3,000 rpm for 30 seconds, and water removed. The sample was then added to the washed hydrazide resin and incubated with end-over-end mixing at room temperature overnight. The resin was centrifuged at 2,500×*g* for 5 min, and the unbound peptides (flow-through/supernatant and the washes) were the non-glycosylated peptides. The resin was then washed three times each with 1.5 M NaCl in water, water, and 100 mM ammonium bicarbonate in water (pH=7.5). Between washes, the centrifugations were at 2,500×*g* for 5 min up to the second water wash, then 2,500×*g* for 6 min up to the first ammonium bicarbonate wash, and then 2,500×*g* for 10 min. Subsequently, the resin was resuspended in 300 µL of G7 reaction buffer (NEB), 30 U/µL of PNGase F (NEB) were added, and incubated at 37°C overnight with shaking. Next, the sample was centrifuged at 2,500×*g* for 10 min and the supernatant (and the below washes) was collected to a triply water-washed 1-dram glass vial. The resin was washed twice with water. Further, the resin was resuspended in 300 µL of 0.2 M citrate-phosphate buffer (pH=5.0), mixed with 12 µL of PNGase A (Roche) and incubated at 37°C overnight with shaking. Next, the sample was centrifuged at 2,500×*g* for 10 min and the supernatant (and the below washes) was collected to the same triply water-washed 1-dram glass vial in which the PNGase F supernatant was. The resin was washed twice with water. The PNGase F and PNGase A-released sample consisted of N-linked glycosylated peptides. To release the O-glycosylated peptides, the resin was incubated with 600 µL of 0.1 M NaOH for 4 hours at 45°C with shaking, 600 µL of 0.3 M acetic acid in water was added on ice, and the supernatant from a 2,500×*g* for 10 min centrifugation was collected. Next, the non-glycan sample was dried to a final volume of around 20% using speed-vac. Further, non-glyco, N-glyco, and O-glyco peptide samples were purified with tC18 sep-pak (Waters) as above, with 0.9 mL of 50% acetonitrile with 0.1% TFA in water elution step. Finally, the samples were dried using speed-vac.

Other protein samples were digested with sequencing grade modified trypsin (Promega Corp) at pH=8.3 (50 mM ammonium bicarbonate) overnight at 37°C with slight shaking. Peptides were acidified with 0.1% TFA and purified using C18 Zip-tips (Millipore) according the manufacturer's instructions and final elutions were dried to around 10 uL. 3-5 uL were loaded onto the LC-MS/MS system.

Peptides were analyzed by positive ion mode liquid chromatography tandem mass spectrometry (LC-MS/MS) using a high resolution hybrid Orbitrap Elite mass spectrometer (Thermo Fisher Scientific) via CID with data-dependent analysis (DDA) using a Top 5 approach (1 full FT-MS scan followed by 5 MS/MS CID scans). Maximum injection time was 50 msec for MS and 100 msec for MS/MS with 1 microscan for both modes. Isolation width was 2.3 Da and dynamic exclusion time was set to 90 sec. Peptides were delivered and separated using an EASY-nLC I nanoflow HPLC (Thermo Fisher Scientific) at 300 nL/min using self-packed 15 cm length × 75 µ m i.d. C18 fritted microcapillary Picofrit columns (New Objective). Solvent gradient conditions were 140 minutes from 3% B buffer to 38% B (B buffer: 100% acetonitrile; A buffer: 0.1% formic acid/99.9% water). MS/MS spectra were analyzed using the Mascot 2.5 search engine by searching the reversed and concatenated UniProt DROME protein database (version 20130918 containing 21,002 protein sequence entries) and the dmel-all translation database (version r522, 2012, 43571 entries) with a parent ion tolerance of 18 ppm and fragment ion tolerance of 0.80 Da. Carbamidomethylation of Cys (+57.0293 Da) was specified in Sequest as a fixed modification and oxidation of Met (+ 15.9949 Da) and deamidation of Asn/Gln (+ 0.9840 Da) as variable modifications. Results were imported into Scaffold 4.0 software (Proteome Software) with a peptide threshold of around 75%, protein threshold of 95%, resulting in a peptide false discovery rate (FDR) of <1.5%.

**Climbing assays**

Climbing assays were performed as previously described (*10, 21, 72*). Each vial had around 10-20 adults (males and females were separated). On the day of the experiment, flies were transferred to an empty vial (without $CO_2$). For climbing assays, flies were tapped to the bottom of the vial and the number of flies able to climb above 6 cm was counted after 10 seconds. After the climbing tests, flies were transferred back to regular food.

Climbing tests with males and females were performed in separate vials and results in **Fig. 3, A to C,** and **Fig. S1I and S17** are presented as averages of across sexes. In **Fig. 3A**, the biological replicates were *n*=4 from 1 experiment (*LPP-Gal4>*control (*Luc-i*)), *n*=6 cumulative across 2 experiments (*LPP-Gal4>*control (*w-i*)), *n*=20 cumulative across 3 experiments (*LPP-Gal4>*control (*attP*)), *n*=12 cumulative across 2 experiments (*LPP-Gal4>CG31326-i-1*), and *n*=10 cumulative across 2 experiments (*LPP-Gal4>CG2145-i-1*). In **Fig. 3B**, the biological replicates were *n*=12 cumulative across 2 experiments (*LPP-Gal4>*control (*v-i*)), *n*=8 cumulative across 2 experiments (*LPP-Gal4>CG31326-i-2*), *n*=8 cumulative across 2 experiments (*LPP-Gal4>CG31326-i-3*), and *n*=4 cumulative across 2 experiments (*LPP-Gal4>CG4332-i-1*). In **Fig. 3C**, the biological replicates were *n*=14 cumulative across 2 experiments (*LPP-Gal4>*control (*w-i*)), *n*=9 from 1 experiment (*LPP-Gal4>*control (*GFP-i*)), *n*=13 cumulative across 2 experiments (*LPP-Gal4>*control (*Luc-i*)), *n*=13 cumulative across 2 experiments (*LPP-Gal4>CG4332-i-2*), and *n*=7 cumulative across 2 experiments (*LPP-Gal4>CG2145-i-2*).

**Immunostaining and confocal microscopy**

Immunostaining and confocal microscopy were performed as previously described (*10, 14, 26*), with modifications. After dissections (see above), thoraxes were fixed in 4% paraformaldehyde (PFA) in PBS at room temperature with gentle shaking for 1 hour. Other tissues were fixed in 4% paraformaldehyde (PFA) in PBS at room temperature with gentle shaking for 30 min. Tissues were then washed four times for 10 minutes each in PBS with 0.1% Triton X-100 (PBST) at room temperature with gentle shaking. Tissues were then blocked with 5% normal goat serum (NGS, Vector Laboratories) in PBST for 30 minutes at room temperature, with gentle shaking. Next, samples were incubated in primary antibody dilutions in PBST overnight at 4°C with gentle shaking in the dark. Primary antibodies and reagents were: 1:200 mouse anti-myc (clone 9E10, Santa Cruz Biotechnology), 1:500 rabbit anti-myc (clone 71D10, Cell Signaling Technology), neat mouse anti-Cnx99A (ER marker; Developmental Studies Hybridoma Bank)(*73*), 1:500 streptavidin-Alexa Fluor (AF) 647 (Invitrogen), 1:200 rat anti-HA (clone 3F10, Roche), 1:2000 rabbit anti-ref(2)P (gift of Andreas Brech) (*74*), 1:100 mouse anti-ubiquitin (clone FK2, Enzo Life Sciences), and 1:100 phalloidin-AF 660 (Invitrogen). Next, samples were washed 4 times for 10 min per wash in PBST at room temperature in the dark, with gentle shaking. Further, tissues were incubated overnight at 4°C in a 1:500 dilution of secondary antibody in PBST, with gentle shaking in the dark. The secondary antibodies were: donkey anti-mouse AF488 (Invitrogen), donkey anti-rabbit AF594 (Invitrogen), donkey anti-rabbit AF555 (Invitrogen), and donkey anti-rat AF594 (Invitrogen). Next, tissues were washed 4 times, 15 minutes each in PBST at room temperature, with gentle shaking in the dark. Finally, tissues were mounted in Vectashield (Vector Laboratories) with two layers of transparent tape on either side to avoid flattening the tissue with the coverslip.

BODIPY (lipids) staining of FB was done as previously described (*75*). After fixation, abdomens were washed 4 times for 10 min each in PBS at room temperature with gentle shaking. Abdomens were then incubated with 1:1,000 DAPI (Molecular Probes) for 30 min at room temperature with gentle shaking, and washed four times for 15 min each in PBS with gentle shaking. Next, abdomens were incubated with 1:1,000 BODIPY493/503 (Invitrogen) in PBS for 30 min at room temperature with gentle shaking, and washed twice in PBS. Abdomens were mounted as described above.

Confocal images were acquired using Zeiss LSM780. For muscles/thoraxes, around 5-6 animals were dissected, and thorax was cut into 2 or 4 fragments. One (occasionally two) confocal image per fragment piece was taken, representing at least 3 animals. Where indicated, maximum intensity projections were generated using ZEN2012 software (Carl Zeiss). Original (non-adjusted) images are shown, unless otherwise indicated. Where indicated, brightness and contrast were adjusted equally in the whole image and equally between channels and groups using ZEN2012 software (Carl Zeiss). In **Fig. 1B** (lower panel) and **Fig. S2, E to H, and M to P**, brightness and contrast were adjusted to 35 and 3.75, respectively. **Fig. 1B** is a maximum intensity projection (MIP) from 2 slices cropped from the blue region of the same sample shown in **Fig. S2, A to D**. In **Fig. S15**, cropped images (original: 134.95 µm × 134.95 µm; cropped: 101.21 µm × 101.21 µm) are shown. This was done for ease of visualization.

**Image Quantification**

The area of protein aggregates was quantified as previously described (*10*). Controls and experimental groups were analyzed in the same way. Representative single confocal slice images (uncropped) of p62/ref(2)P staining were converted to single color tiff files using ZEN 2012 (Carl Zeiss). Next, using FIJI (FIJI is just image J) (*76*), images were converted to 8-bit grayscale, and threshold was adjusted in order to use the analyze particles function. The maximum entropy method (*77*) was used in **Fig. 3, D, E, G, and H** and the triangle method (*78*) was used in **Fig. 3F**. Next, the analyze particles function was used, in which the minimum particle size was 10 pixels$^2$ in **Fig. 3, D, E, G, and H**, and 20 pixels$^2$ in **Fig. 3F**. Aggregate areas were normalized to controls.

**Data analysis and statistics**

Data was analyzed using ZEN2012 software (Carl Zeiss), FIJI, Microsoft Excel, Graphpad Prism 7, and OriginPro 2017. Data is shown as mean ± standard error of the mean (SEM). Biological replicates are shown, unless otherwise indicated.In **Fig. 1C**, using Graphpad Prism, in the top panel, a one-way ANOVA was performed, together with an unpaired two-tailed t-test with Holm-Sidak correction. In the bottom panel, an unpaired two-tailed t-test was performed. In **Fig. 2, B to F**, a two-sided chi-square test was performed using Graphpad Prism. In **Fig. 3, A to C**, using Graphpad Prism, a one-way ANOVA was performed, together with Linear Two-Stage Step-Up false-discovery rate (FDR) calculation of Benjamini, Krieger, Yekutieli. Please note that for each RNAi stock center collection, to account for multiple comparisons testing, the statistical calculation took into account additional tested *LPP-Gal4>RNAi* groups which did not show a phenotype in both climbing-ability and protein aggregate formation assays (data not shown). Adjusted *p*-values (*q*-values) are shown. In **Fig. 3, D to H**, an unpaired two-tailed t-test (assuming unequal variances for h) using Microsoft Excel.

In **Fig. S1, A to H**, log-rank test using the Mantel-Cox procedure was performed using Graphpad Prism. In **Fig. S1I**, using Graphpad Prism, a one-way ANOVA was performed, together with Linear Two-Stage Step-Up false-discovery rate (FDR) calculation of Benjamini, Krieger, Yekutieli. Adjusted *p*-values (*q*-values) are shown. In **Fig. S6C**, a linear regression analysis was performed using Graphpad Prism. In **Fig. S6D; S7, B and C**; and **S8**, using Graphpad Prism, a two-sided chi-square test was performed. In **Fig. S9A**, using Graphpad Prism, a one-way ANOVA and Kolmogorov-Smirnov tests were performed. In **Fig. S9C**, using Graphpad Prism, a two-sided chi-square test was performed. In **Fig. S9D**, OriginPro 2017 was used for the statistical calculation and data was graphed using Graphpad Prism. A bigaussian fit was performed on the data, and the fitted peak centers ($x_c$) were compared using the F-test and Aikaike's Information Criterion Test (AIC). In **Fig. S10, C, and E to H**, using Graphpad Prism, a two-sided chi-square test was performed. In **Fig. S10, I and J**, using Graphpad Prism, a chi-square test for trend was performed. In **Fig. S11B**, using Graphpad Prism, a two-sided chi-square test was performed between observed and expected values. In **Fig. S13**, using Graphpad Prism, a two-sided chi-square test was performed. In **Fig. S17**, using Graphpad Prism, a one-way ANOVA was performed, together with Linear Two-Stage Step-Up false-discovery rate (FDR) calculation of Benjamini, Krieger, Yekutieli. Please note that for each RNAi stock center collection, to account for multiple comparisons testing, the statistical calculation took into account additional tested *DMEF2-Gal4>RNAi* groups which did not show a phenotype in both climbing-ability and protein aggregate formation assays (data not shown). Adjusted *p*-values (*q*-values) are shown.

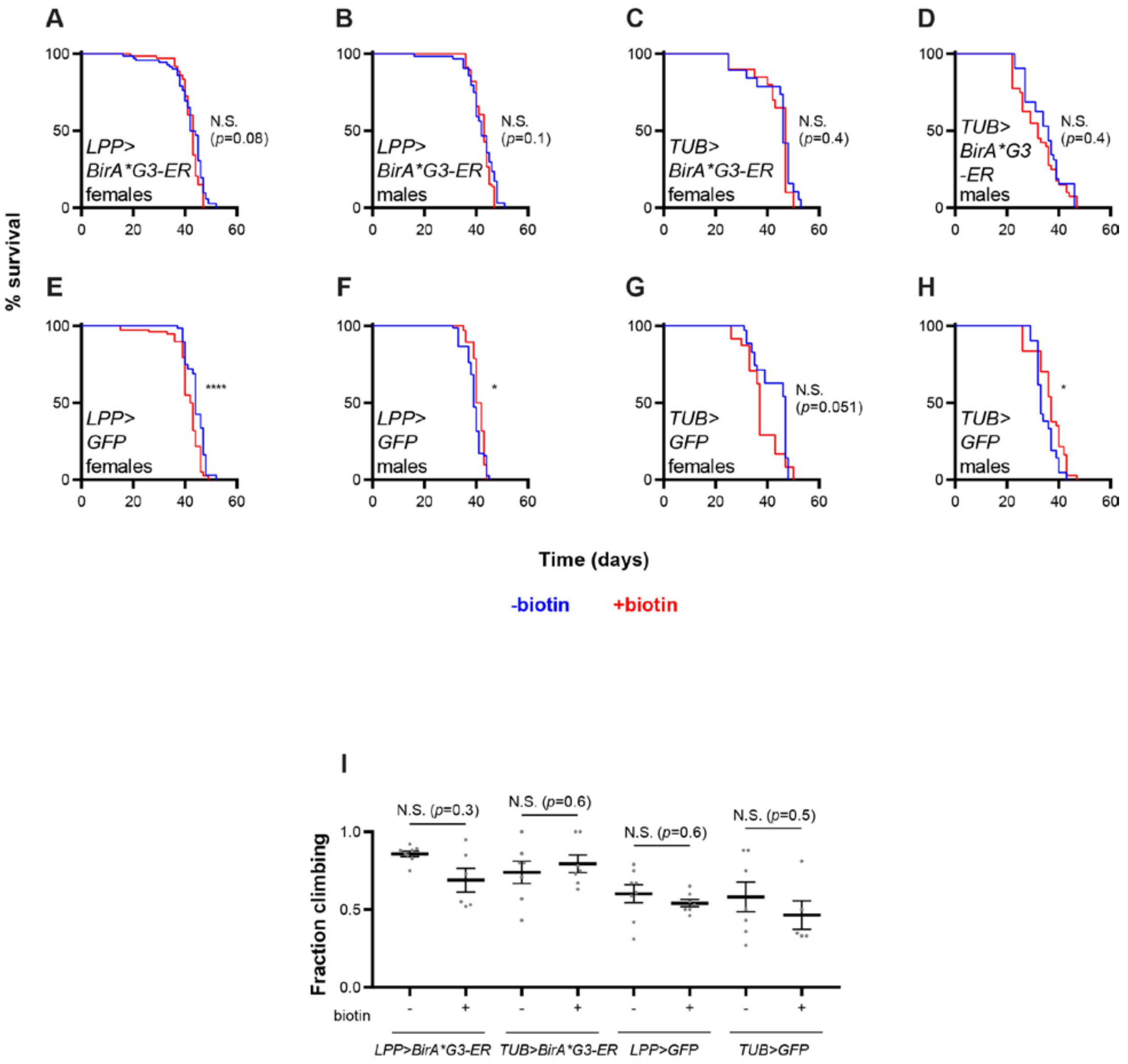


**Fig. S1: Whole body or fat body (FB) biotinylation does not significantly affect adult fly survival or climbing-ability (Fig. 1 supplement).**

Adults were grown on regular food and switched to regular food (-biotin, blue) or biotin-containing food (+biotin, red) at 29°C.

**(A to H)** Statistics: Log-rank test. N.S. means not significant. **(A)** *LPP-Gal4>BirA*G3-ER* females. –biotin: *n*=72 flies, median lifespan=43 days. +biotin: *n*=73 flies, median lifespan=43 days. N.S. (*p*=0.081). **(B)** *LPP-Gal4>BirA*G3-ER* males. –biotin: *n*=64 flies, median lifespan=42 days. +biotin: *n*=79 flies, median lifespan=43 days. N.S. (*p*=0.1035). **(C)** *TUB-Gal4>BirA*G3-ER* females. –biotin: *n*=19 flies, median lifespan=46 days. +biotin: *n*=20 flies, median lifespan=47 days. N.S. (*p*=0.43). **(D)** *TUB-Gal4>BirA*G3-ER* males. –biotin: *n*=32 flies, median lifespan=36 days. +biotin: *n*=40 flies, median lifespan=32 days. N.S. (*p*=0.42). **(E)** *LPP-Gal4>mCD8-GFP* females. –biotin: *n*=68 flies, median lifespan=44 days. +biotin: *n*=78 flies, median lifespan=42.5 days. ****$p<0.0001$. **(F)** *LPP-Gal4>mCD8-GFP* males. –biotin: *n*=76 flies, median lifespan=39 days. +biotin: *n*=104 flies, median lifespan=41 days. *$p$=0.015. **(G)** *TUB-Gal4>mCD8-GFP* females. –biotin: *n*=35 flies, median lifespan=47 days. +biotin: *n*=24 flies, median lifespan=37 days. N.S. (*p*=0.051). **(H)** *TUB-Gal4>mCD8-GFP* males. –biotin: *n*=21 flies, median lifespan=33 days. +biotin: *n*=37 flies, median lifespan=37 days. *$p$=0.036.

**(I)** Climbing-ability assays in 3 week old flies at 29°C. Biological replicates: *n*=8 (*LPP-Gal4>BirA*G3-ER* –biotin), *n*=6 (*LPP-Gal4>BirA*G3-ER* +biotin), *n*=7 (*TUB-Gal4>BirA*G3-ER* –biotin), *n*=7 (*TUB-Gal4>BirA*G3-ER* +biotin), *n*=8 (*LPP-Gal4>mCD8-GFP* –biotin), *n*=7 (*LPP-Gal4>mCD8-GFP* +biotin), *n*=7 (*TUB-Gal4>mCD8-GFP* –biotin), *n*=5 (*TUB-Gal4>mCD8-GFP* +biotin). Statistics: mean±SEM; one-way ANOVA and Benjamini, Krieger, Yekutieli Linear Two-Stage Step-Up FDR.

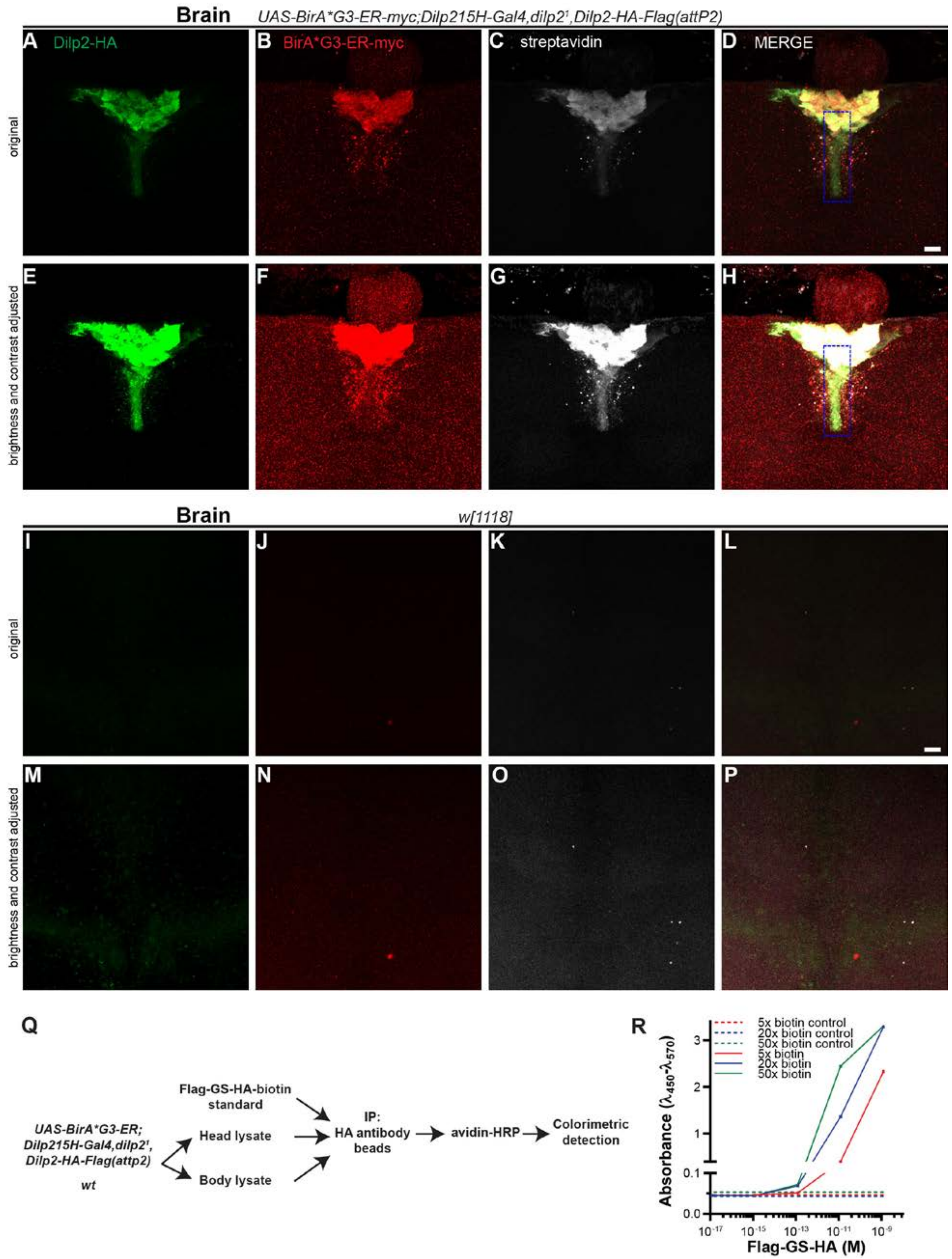


**Fig. S2: Biotinylated Dilp2 (*Drosophila* insulin-like peptide 2) detection (Fig. 1 supplement).**

**(A to P)** *UAS-BirA*G3-ER-myc;Dilp215H-Gal4,dilp2$^1$,Dilp2-HA-Flag(attP2)* **(A to H)** or *w[1118] wt* control **(I to P)** brains were stained for HA (green), myc (red), and streptavidin (white). Representative maximum intensity projections are shown. **(A to D)** and **(I to L)** are original images, and **(E to H)** and **(M to P)** are images with equally-adjusted brightness and contrast. In **(D)** and **(H)**, the blue rectangle is the region from which **Fig. 1B** (2 confocal slices) was cropped. Scale bar: 10 µm. Flies were fed with 50 µM biotin during adulthood.

**(Q)** Procedure for spatially informed bead enzyme-linked immunosorbent assay (ELISA) of biotinylated Dilp2-HA.

**(R)** Calibration curve of synthesized Flag-GS-HA standard peptide, biotinylated with different molar excess of sulfo-NHS-biotin. Shown are means±SEM of three repeat spectrophotometric measurements of the same sample. Dashed lines are absorbances of negative control samples (PBS treated with sulfo-NHS-biotin).

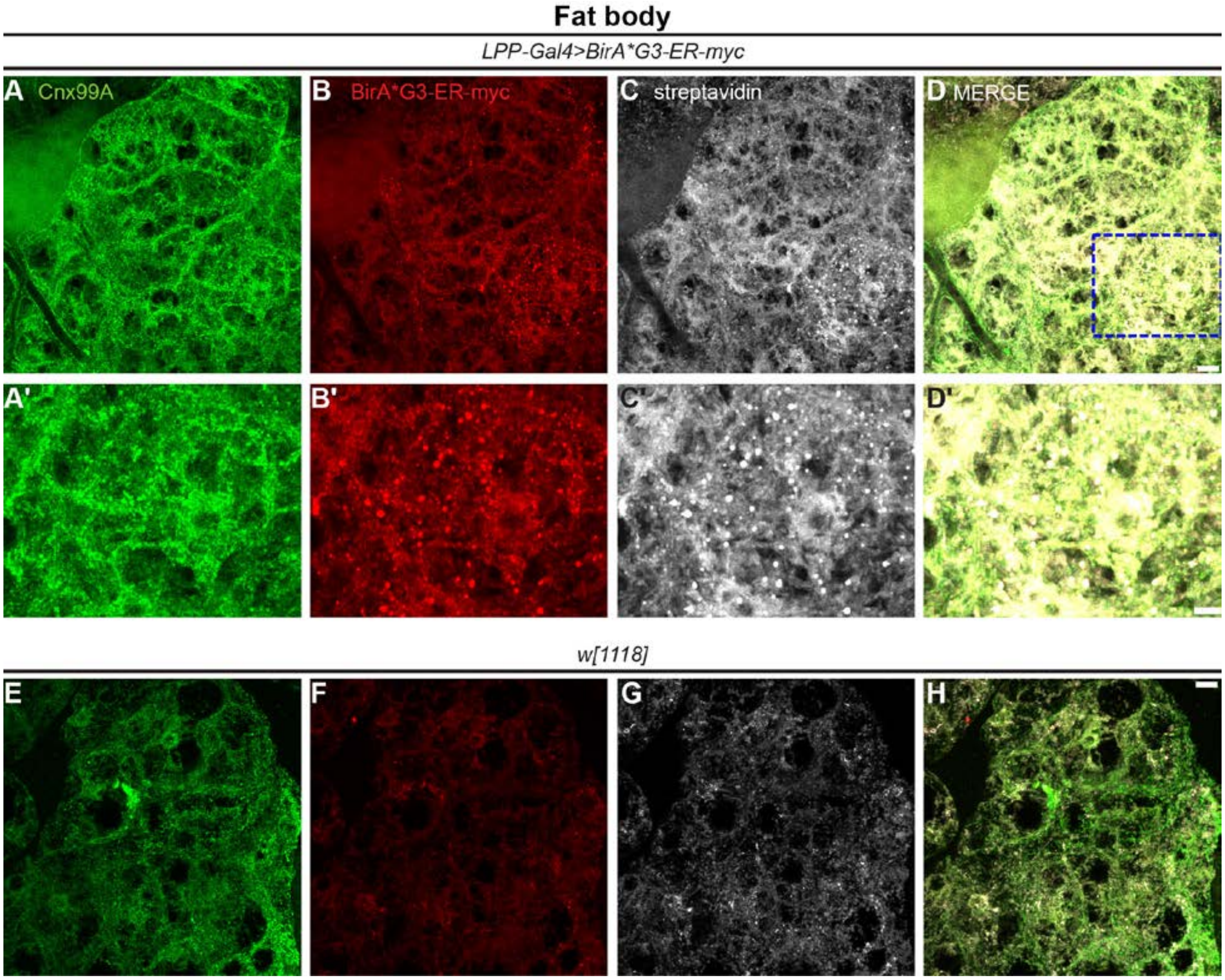


**Fig. S3: Detection of BirA*G3-ER and biotin in the adult abdominal fat body (FB) (Fig. 1 supplement).**

*LPP-Gal4>UAS-BirA*G3-ER-myc* **(A to D)** and *w[1118] wt* control **(E to H)** samples were stained for ER marker Cnx99A(*73*) (green), myc (red), and streptavidin (white). Representative maximum intensity projections are shown. **(A' to D')** are zoomed in images from blue rectangle in **(A to D)**. Scale bar: 10 µm **(A to D, E to H)**; 5 µm **(A' to D')**. Flies were fed with 50 µM biotin during adulthood.

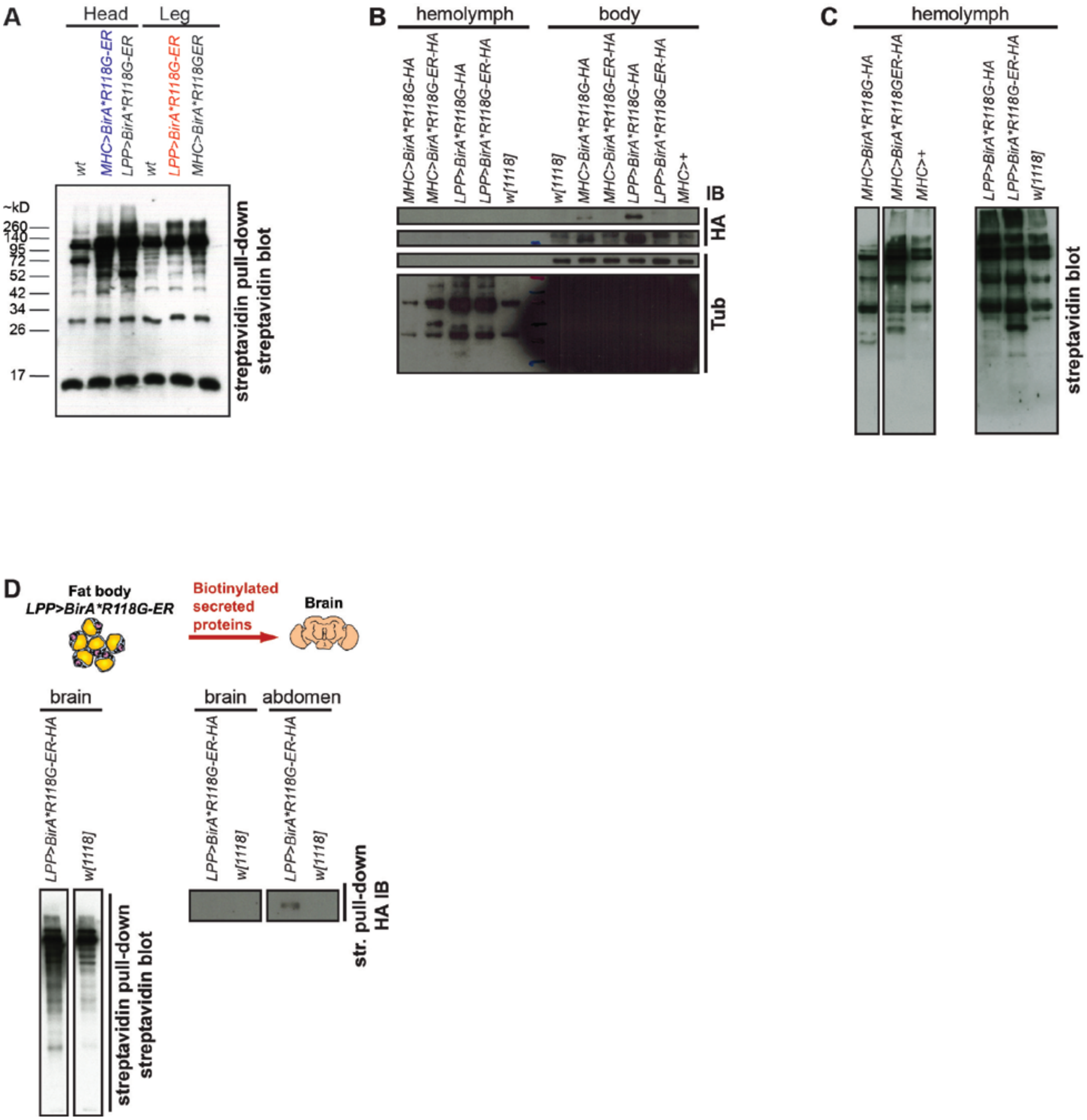


**Fig. S4: Organ-of origin biotinylation using BirA*R118G and detection of biotinylated proteins in distal organs or body parts. (Fig. 1 supplement).**

**(A)** Streptavidin bead pull-down followed by streptavidin-HRP detection in head and leg lysates after muscle (*MHC-Gal4*) and fat body (FB, *LPP-Gal4*) biotinylation using BirA*R118G-ER. Genotypes (males and females): *w[1118] wt*, *UAS-BirA*R118G-ER(attP40)/UAS-BirA*R118G-ER(attP40);MHC-Gal4/MHC-Gal4* and *UAS-BirA*R118G-ER(attP40)/UAS-BirA*R118G-ER(attP40);LPP-Gal4/UAS-BirA*R118G-ER(attP2)*.

**(B)** BirA*R118G-HA or BirA*R118G-ER-HA is detected in body but not hemolymph (blood). For HA and Tubulin (Tub), lower (top panel) and higher (lower panel) exposures are shown. Note that the body BirA*R118G-ER-HA is detected at lower levels than BirA*R118G (cytoplasmic/nuclear); however, this signal is consistently detected in multiple experiments. Genotypes (females): *w[1118] wt*, *UAS-BirA*R118G(attp40)/UAS-BirA*R118G(attP40);MHC-Gal4/MHC-Gal4*, *UAS-BirA*R118G-ER(attP40)/UAS-BirA*R118G-ER(attP40);MHC-Gal4/MHC-Gal4, UAS-BirA*R118G(attP40)/UAS-BirA*R118G(attP40);LPP-Gal4/LPP-Gal4*, *UAS-BirA*R118G-ER(attP40)/UAS-BirA*R118G-ER(attP40);LPP-Gal4/LPP-Gal4*, *+/+;MHC-Gal4*. IB means immunoblot.

**(C)** Detection of biotinylated proteins in the hemolymph after muscle and FB biotinylation using BirA*R118G-ER and BirA*R118G (cytoplasmic/nuclear biotinylation for unconventional protein secretion). Genotypes (females): *w[1118] wt*, *UAS-BirA*R118G(attP40)/UAS-BirA*R118G(attP40);MHC-Gal4/MHC-Gal4*, *UAS-BirA*R118G-ER(attP40)/UAS-BirA*R118G-ER(attP40);MHC-Gal4/MHC-Gal4, UAS-BirA*R118G(attP40)/UAS-BirA*R118G(attP40);LPP-Gal4/LPP-Gal4*, *UAS-BirA*R118G-ER(attP40)/UAS-BirA*R118G-ER(attP40);LPP-Gal4/LPP-Gal4*, *+/+;MHC-Gal4*.

**(D)** Abdomens and FB-free brains were dissected from *w[1118] wt* and *UAS-BirA*R118G-ER(attP40)/UAS-BirA*R118G-ER(attP40);LPP-Gal4/UAS-BirA*R118G-ER(attP2)* male adult flies. Blots for streptavidin-HRP (left) and HA (right) were performed. Extra biotinylated proteins were detected in brains of *LPP>BirA*R118G-ER* flies, while HA is detected in abdomens of *LPP>BirA*R118G-ER* flies only.

In all panels, flies were maintained with 50 µm biotin in food during adulthood. *wt* means *wild-type*.

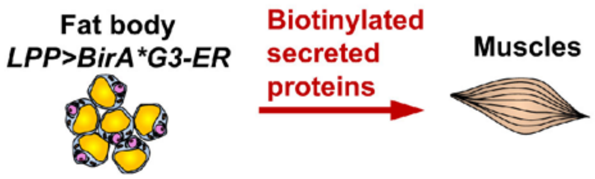


| Sample | TMT label |
|---|---|
| *LPP>BirA*G3-ER* legs | 127N |
| *LPP>BirA*G3-ER* legs | 129N |
| *wt* legs | 126 |
| *wt* legs | 128C |
| *MHC>BirA*G3-ER* heads | 128N |
| *MHC>BirA*G3-ER* heads | 130N |
| *wt* heads | 127C |
| *wt* heads | 129C |

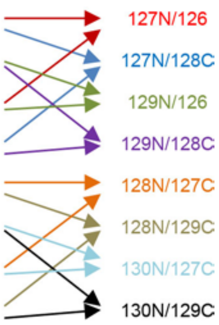


**Fig. S5: Experimental setup for the identification of fat body (FB)-derived proteins in legs and muscle-derived proteins in heads in a single quantitative tandem mass-tag (TMT) mass spectrometry (MS) experiment (Fig. 2 supplement).**

The different TMT state signals were compared to generate TMT ratios (right of the arrows). There were 4 TMT ratio comparisons for legs and 4 for heads. Genotypes: *w[1118] wt (wild-type)*, *LPP-Gal4>UAS-BirA*G3-ER*, *MHC-Gal4>UAS-BirA*G3-ER*. Flies were maintained with 50 µm biotin in food during adulthood.

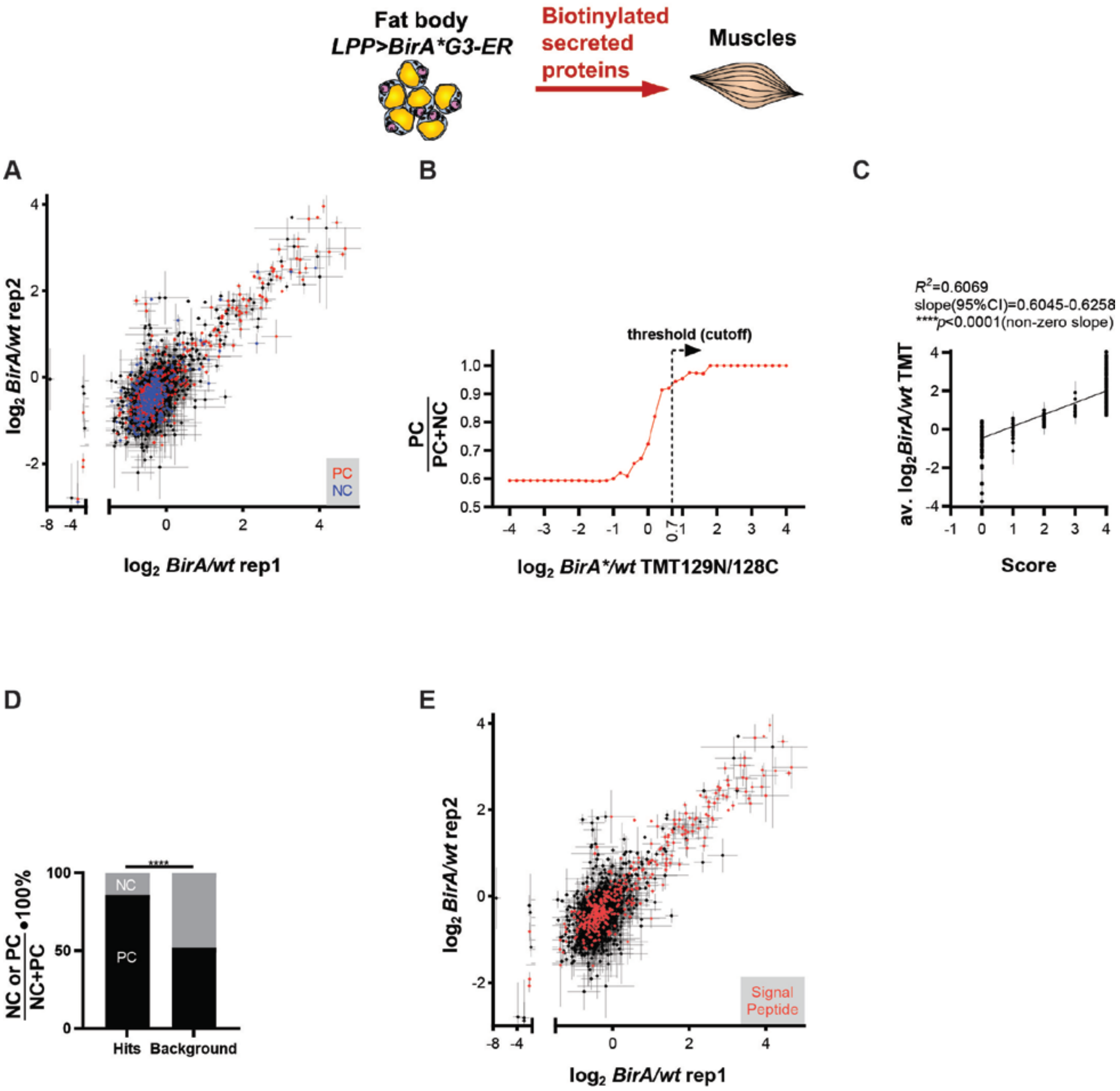


**Fig. S6: Thresholding and signal peptide analyses of biotinylated leg/muscle proteins originating from fat body (FB) tandem mass-tag (TMT) mass spectrometry (MS) using BirA*G3-ER labeling (Fig. 2 supplement).**

Genotypes: *w[1118] wt (wild-type)*, *LPP-Gal4>UAS-BirA*G3-ER*. Flies were maintained with 50 µm biotin in food during adulthood.

**(A)** Leg $log_2$(*BirA*G3-ER / wt*) TMT-ratios in two replicates, with each point is *n*=2 comparisons, mean±SEM $log_2$TMT ratio. Proteins identified with MS were compared to positive control (PC) secreted protein/receptor (red points) and negative control (NC, intracellular) (blue points).

**(B)** Representative $\frac{\#PC}{\#PC+\#NC}$ versus *BirA*G3-ER/wt* TMT ratio graph (out of four). The threshold TMT ratio was chosen at which $\frac{\#PC}{\#PC+\#NC} > 0.9$.

**(C)** Increased TMT-ratios are associated with higher MS scores. Each point is a mean±SEM $\log_2$TMT ratio for each identified protein. Linear regression results are presented.

**(D)** Hits (score≥1) have higher $\frac{\#PC}{\#PC+\#NC}$. ****$p$<0.0001.

**(E)** Proteins with an identified signal peptide (*36*) (red) were mapped onto the leg $\log_2$(*BirA*G3-ER/wt*) TMT-ratios in two replicates graph. Each point is $n$=2 comparisons, mean±SEM.

Statistics **(D)**: Chi-square test.

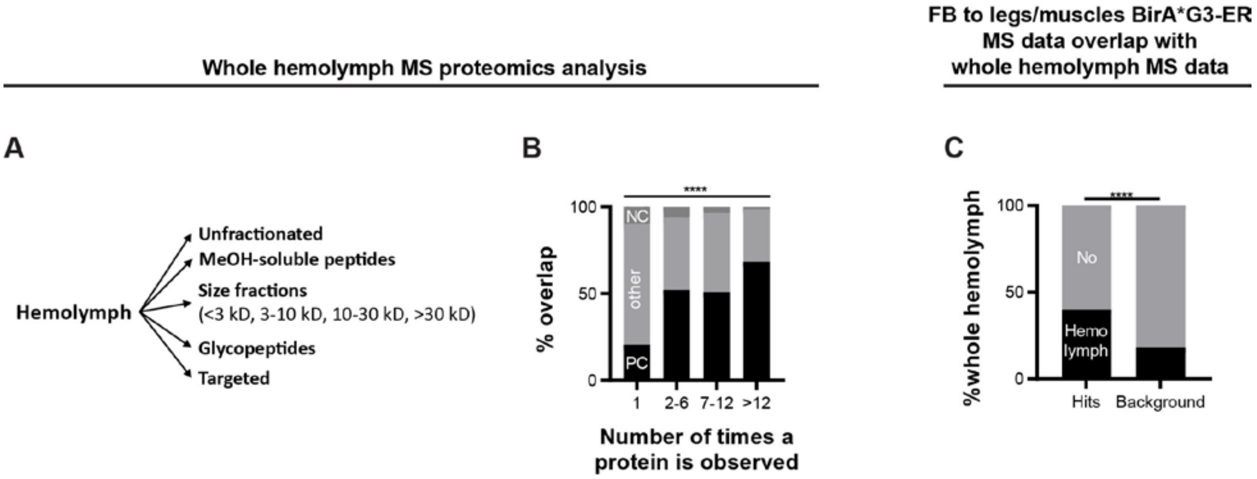


**Fig. S7: Whole hemolymph MS proteomics analysis of legs/muscles proteins derived from fat body (FB) identified in the BirA*G3-ER MS dataset (Fig. 2 supplement).**

**(A)** Summary of the hemolymph processing in total hemolymph MS proteomics experiments (see **Materials and Methods** section for more details).

**(B)** Each identified protein in total fly hemolymph MS was assigned to a positive control (PC) secreted/receptor, negative control (NC) intracellular, or other (unknown) categories (see **Materials and Methods**). The x-axis is the number of times a protein is observed (or sum of unique peptides across all experiments). We identified a total of 1561 proteins, including 688 proteins of poorly-characterized functions (“Computed Genes”/CGs; FlyBase (*42*) r551). ****$p$<0.0001.

**(C)** Hits (score≥1) from the legs/muscles proteins derived from FB (identified in the BirA*G3-ER MS dataset) are enriched for proteins identified in whole fly hemolymph. ****$p$<0.0001.

Statistics **(B and C)**: Chi-square test.

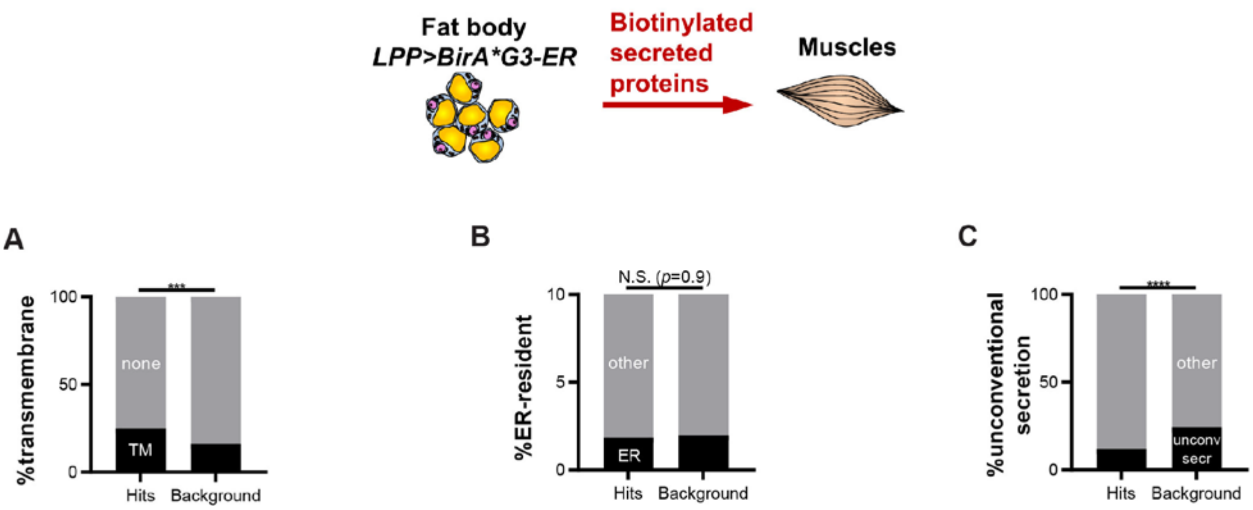


**Fig. S8: Transmembrane, ER-resident, and unconventional secretion analyses of fat body (FB)-derived proteins present in legs/muscles, as identified using tandem mass-tag (TMT) mass spectrometry (MS) using BirA*G3-ER labeling (Fig. 2 supplement).**

**(A)** Hits (score≥1) are enriched for proteins with transmembrane domains (*37*). ****p*=0.0005.

**(B)** Hits (score≥1) are not enriched for ER-resident proteins.

**(C)** Hits (score≥1) are de-enriched for proteins predicted to be secreted unconventionally(*38*).

*****p*<0.0001.

Statistics: Chi-square test.

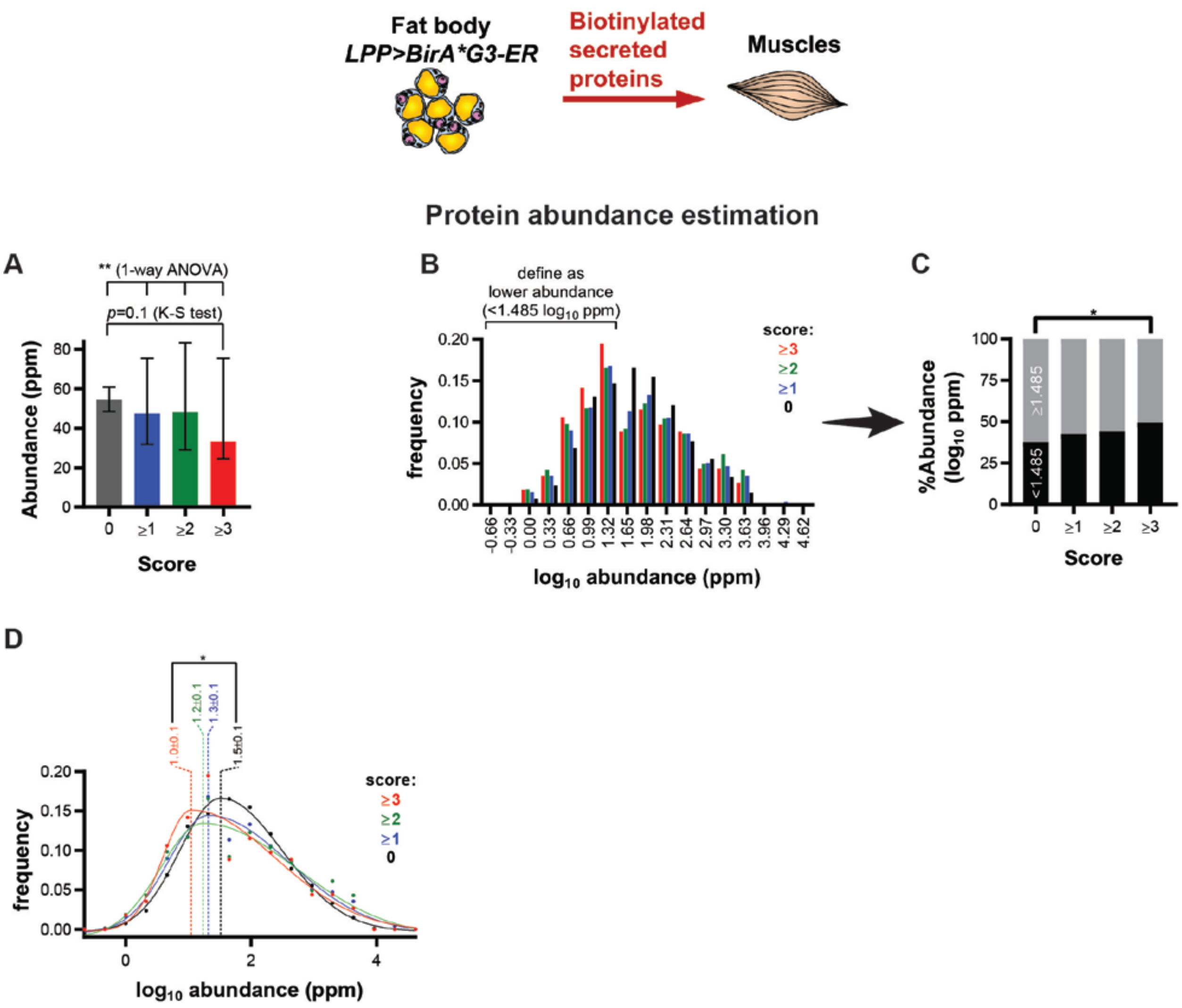


**Fig. S9: BirA*G3-ER legs/muscles FB-derived hits were enriched for lower-abundance proteins (Fig. 2 supplement).**

Protein abundance information was from integrated entire organism PAX database for *Drosophila melanogaster* (*41*).

**(A)** Increased mass spectrometry (MS) scores show a trend for decreased protein abundance. $^{**}p$=0.0033 (one-way ANOVA between all groups); *p*=0.10 (Kolmogorov-Smirnov (K-S) test between score=0 and score≥3).

**(B)** Frequency vs $\log_{10}$ protein abundance plot.

**(C)** Histogram of abundance versus score, when lower abundance is <1.485 $\log_{10}$ ppm, reveals a general trend towards lower protein abundances at higher MS scores. Chi-square test: *$p$=0.013 between MS score=0 and MS score≥3.

**(D)** Frequency vs $\log_{10}$ protein abundance plot with bigaussian fits. Score=0: $x_c$ (peak center±SEM)=1.5±0.1 $\log_{10}$ppm; $R^2$=0.9925. Score≥1 $x_c$=1.3±0.1 $\log_{10}$ppm; $R^2$=0.96039. Score≥2 $x_c$=1.2±0.1 $\log_{10}$ppm; $R^2$=0.91765. Score≥3 $x_c$=1.0±0.1 $\log_{10}$ppm; $R^2$=0.90396. $x_c$(score=0) versus $x_c$(score≥3): F-test *$p$=0.032; AIC (Akaike's Information Criterion Test) weights suggest that $x_c$ are different (same=0.24884, different=0.75116).

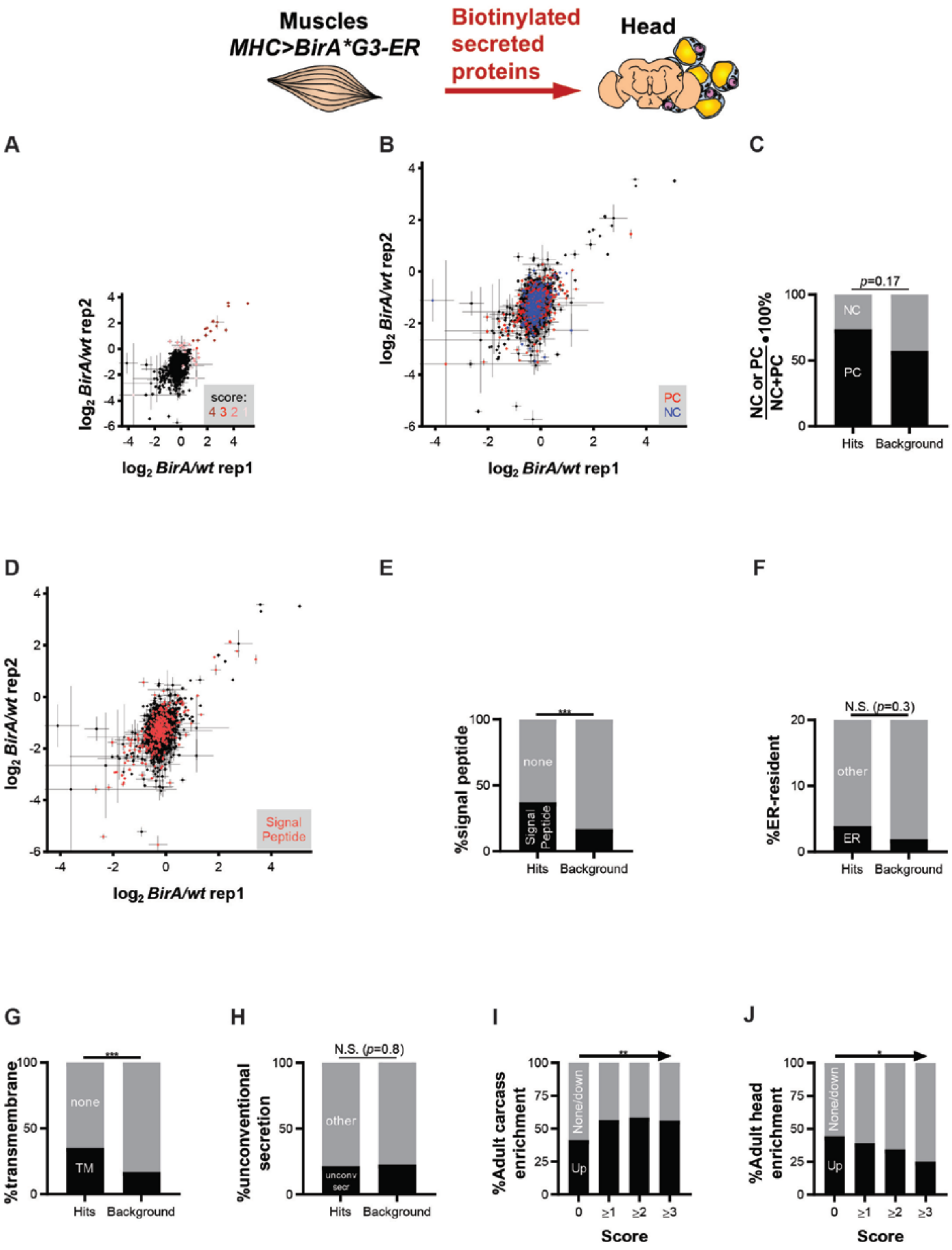


**Fig. S10: Identification of muscle proteins present in heads using BirA*G3-ER (Fig. 2 supplement).**

Muscles were labeled using *MHC-Gal4>BirA*G3-ER* (endoplasmic reticulum) and biotinylated proteins from heads were analyzed using tandem mass-tag (TMT) mass spectrometry (MS). *w[1118] wt (wild-type)* heads were used as controls. Flies were maintained with 50 µm biotin in food during adulthood.

**(A)** Head $\log_2$(*BirA*G3-ER/wt*) TMT ratios in two replicates. Each point is *n*=2 comparisons, mean±SEM $\log_2$TMT ratio. MS score: number of comparisons (from 4) in which TMT ratio>threshold (score 4 is for most confident hits and 0 is background). For each of the four *BirA*G3-ER/wt* TMT-ratio comparisons, we determined threshold TMT ratios for hit-calling as described in the **Materials and Methods** section. A total of 51 muscle endoplasmic reticulum (ER)-derived proteins targeting heads were identified.

**(B)** Head proteins identified with MS were compared to positive control (PC) secreted protein/receptor (red points) and intracellular negative control (NC) (blue points).

**(C)** Hits (score≥1) trend towards higher $\frac{\#PC}{\#PC+\#NC}$.

**(D)** Proteins with an identified signal peptide(*36*) (red) were mapped onto the head $\log_2$(*BirA*G3-ER/wt*) TMT-ratios in two replicates graph. Each point is *n*=2 comparisons, mean±SEM $\log_2$TMT ratio.

**(E)** Hits (score≥1) have a higher fraction of proteins with putative signal peptides(*79*). ****p*=0.0001.

**(F)** Hits (score≥1) are not statistically-significantly enriched for ER-resident proteins.

**(G)** Hits (score≥1) are enriched for proteins with transmembrane domains(*37*). ****p*=0.0007.

**(H)** Hits (score≥1) are not enriched for proteins predicted to be secreted unconventionally(*38*).

Statistics **(C, E to H)**: Chi-square test.

**(I to J)** Higher MS scores correlate with a higher fraction of proteins enriched for adult carcass (which includes muscles) mRNA microarray(*39*) **(I)**, and with a lower fraction of proteins enriched for adult head mRNA microarray(*39*) **(J)**. Chi-square test for trend: ***p*=0.0063, **p*=0.043.

### Comparison between fat body to legs and muscles to heads datasets

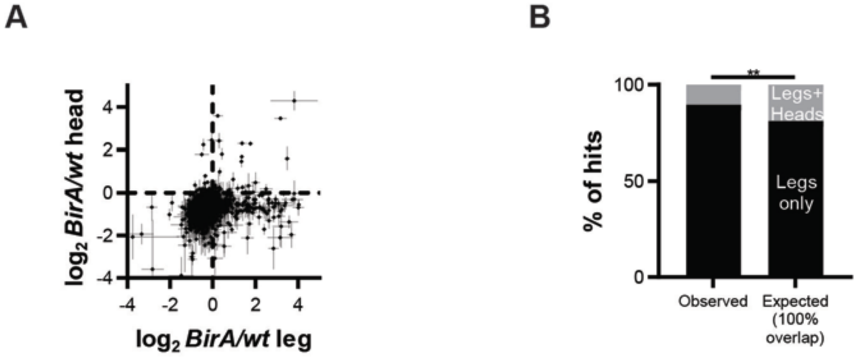


**Fig. S11: Comparison between BirA*G3-ER legs and heads datasets reveals limited overlap, suggestive of organ-specificity in secretome trafficking (Fig. 2 supplement).**

**(A)** Head $\log_2$(*BirA*G3-ER/wt*) TMT ratios vs leg $\log_2$(*BirA*G3-ER/wt*) TMT ratios. Each point is *n*=4 comparisons; mean±SEM $\log_2$TMT ratio.

**(B)** Comparison between observed and expected (based on 100% overlap) overlap between legs and heads datasets. The observed overlap is less than expected. Chi-square test: ***p*=0.0051.

*wt* means *wild-type*.

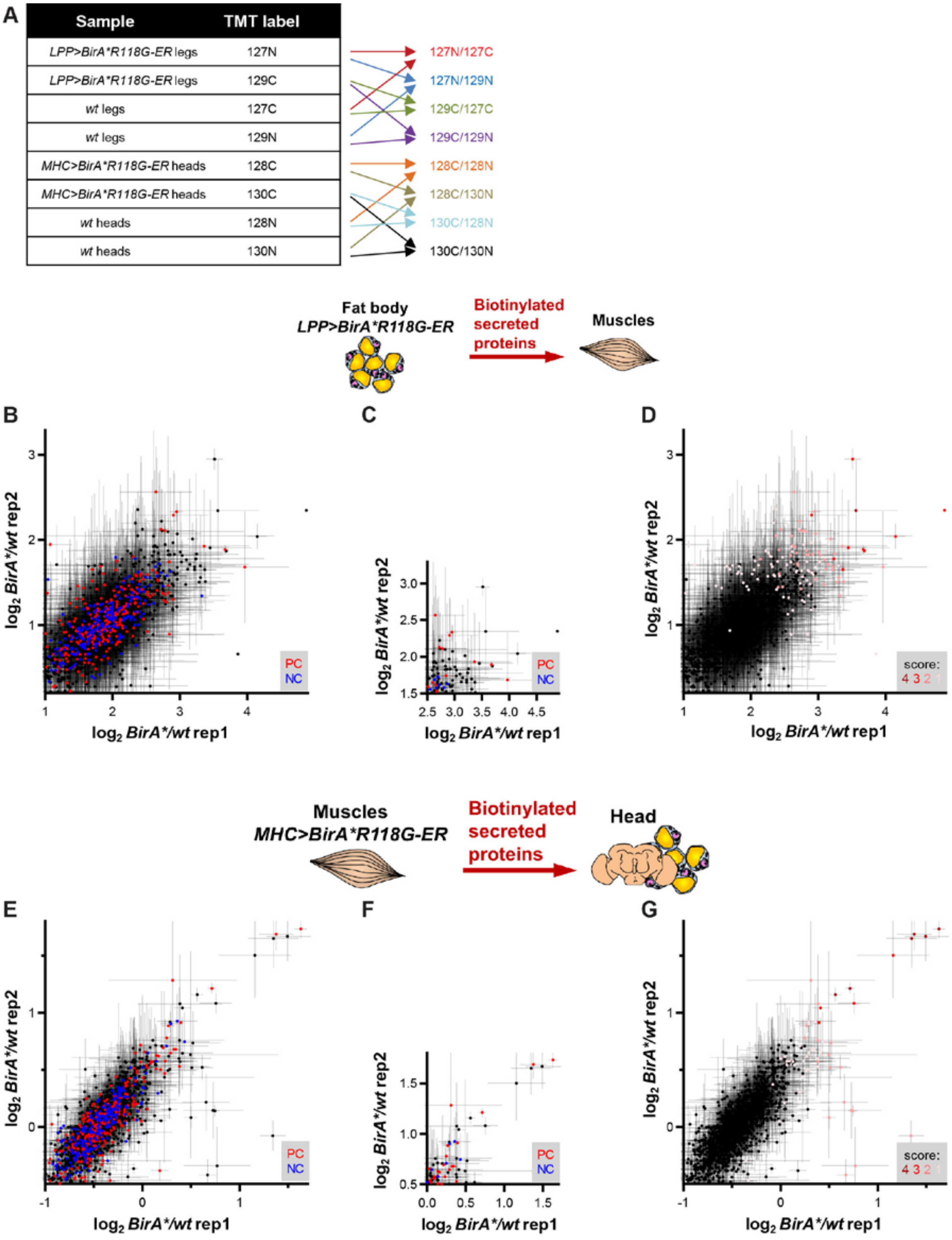


**Fig. S12: Using BirA*R118G-ER to identify legs/muscles proteins produced from fat body (FB) and head proteins produced from muscles (Fig. 2 supplement).**

**(A)** Experimental design. The different TMT state signals were compared to generate TMT ratios (right of the arrows). There were 4 TMT ratio comparisons for legs and 4 for heads. Genotypes: *w[1118] wt (wild-type)*, *UAS-BirA*R118G-ER(attP40)/UAS-BirA*R118G-ER(attP40);MHC-Gal4/MHC-Gal4* and *UAS-BirA*R118G-ER(attP40)/UAS-BirA*R118G-ER(attP40);LPP-Gal4/UAS-BirA*R118G-ER(attP2)*. Flies were maintained with 50 µm biotin in food during adulthood.

**(B to D)** Identification of legs/muscles proteins produced in FB. **(B)** Leg $log_2$(*BirA*R118G-ER/wt*) TMT ratios in two replicates. Each point is *n*=2 comparisons, mean±SEM $log_2$TMT ratio. Leg proteins identified with MS were compared to positive control (PC) secreted protein/receptor (red points) and intracellular negative control (NC) (blue points). **(C)** Zoom of graph in (b). **(D)** Leg $log_2$(*BirA*R118G-ER/wt*) TMT ratios in two replicates. Each point is *n*=2 comparisons, mean±SEM $log_2$TMT ratio. MS score: number of comparisons (from 4) in which TMT ratio>threshold (score 4 is for most confident hits and 0 is background). For each of the four *BirA*R118G-ER/wt* TMT-ratio comparisons, we determined threshold TMT ratios for hit-calling, as described in the **Materials and Methods** section.

**(E to G)** Identification of head proteins produced in muscles. **(E)** Head $log_2$(*BirA*R118G-ER/wt*) TMT ratios in two replicates. Each point is *n*=2 comparisons, mean±SEM $log_2$TMT ratio. Head proteins identified with MS were compared to PC and NC lists. **(F)** Zoom of graph in (e). **(G)** Head $log_2$(*BirA*R118G-ER/wt*) TMT ratios in two replicates. Each point is *n*=2 comparisons, mean±SEM $log_2$TMT ratio. MS score: number of comparisons (from 4) in which TMT ratio>threshold (score 4 is for most confident hits and 0 is background). For each of the four *BirA*R118G-ER/wt* TMT-ratio comparisons, we determined threshold TMT ratios for hit-calling, as described in the **Materials and Methods** section.

### Comparison between BirA*G3-ER and BirA*R118G-ER fat body to legs datasets

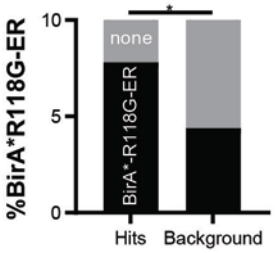


**Fig. S13: Legs *LPP-Gal4>BirA*G3-ER* dataset hits (score ≥ 1) show statistically-significant overlap with *LPP-Gal4>BirA*R118G-ER* dataset hits, compared to background (Fig. 2 supplement).**

Chi-square test: *$p$=0.0144

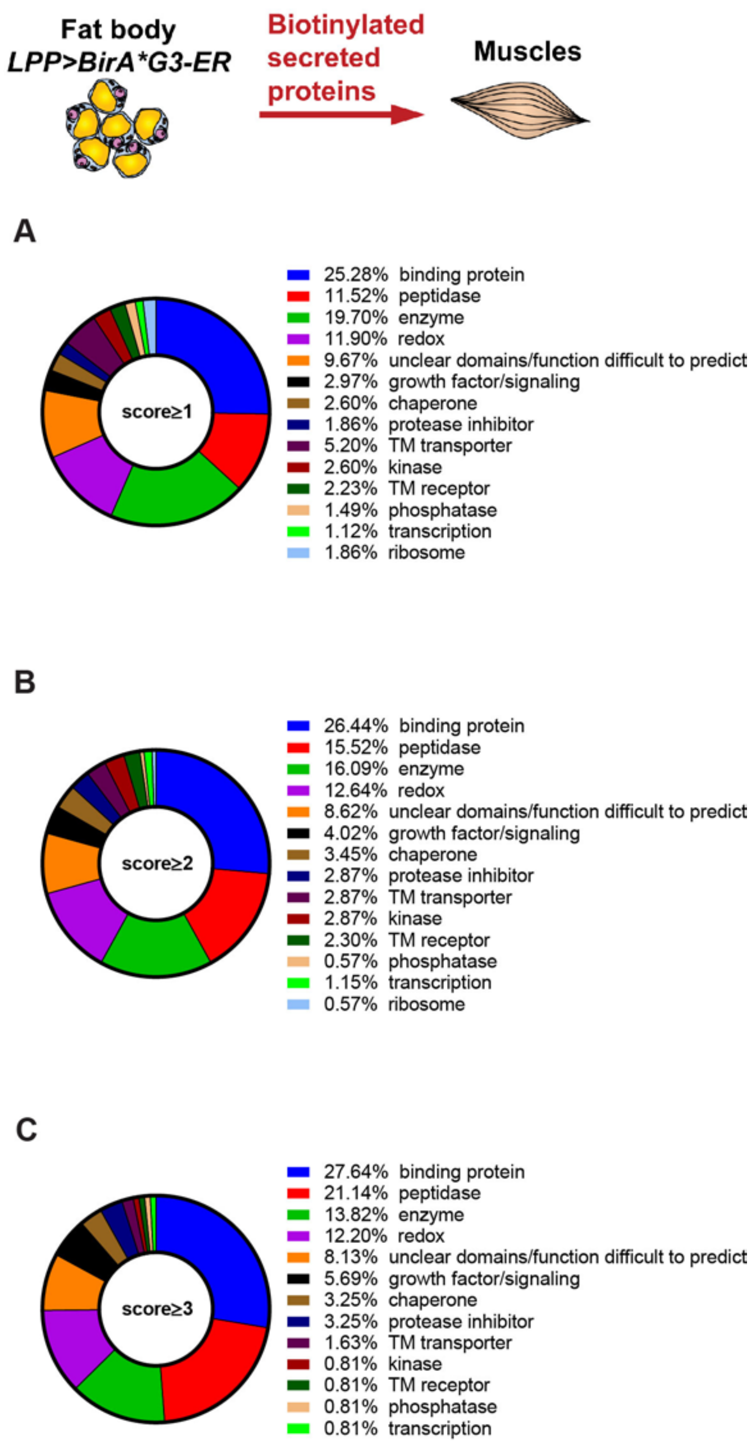


**Fig. S14: Manual categorization of hits (score ≥ 1) according to data from FlyBase (*42*) and ortholog data from NCBI Gene (*43*) (Fig. 2 supplement).**

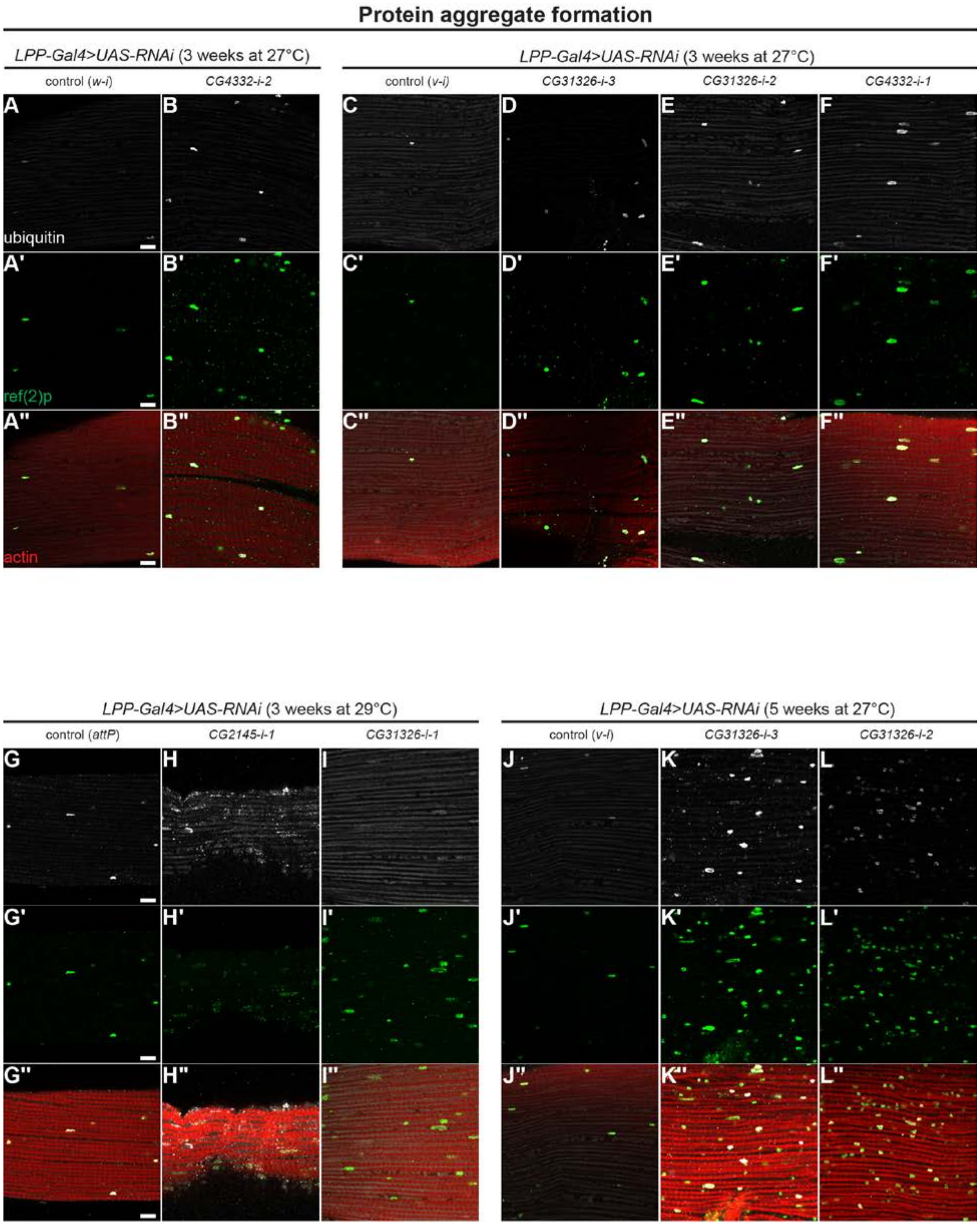
Protein aggregate formation
LPP-Gal4>UAS-RNAi (3 weeks at 27°C)
control (w-i)
CG4332-i-2
LPP-Gal4>UAS-RNAi (3 weeks at 27°C)
control (v-i)
CG31326-i-3
CG31326-i-2
CG4332-i-1
A
B
C
D
E
F
ubiquitin
A'
B'
C'
D'
E'
F'
ref(2)p
A''
B''
C''
D''
E''
F''
actin
LPP-Gal4>UAS-RNAi (3 weeks at 29°C)
control (attP)
CG2145-i-1
CG31326-i-1
LPP-Gal4>UAS-RNAi (5 weeks at 27°C)
control (v-i)
CG31326-i-3
CG31326-i-2
G
H
I
J
K
L
G'
H'
I'
J'
K'
L'
G''
H''
I''
J''
K''
L''

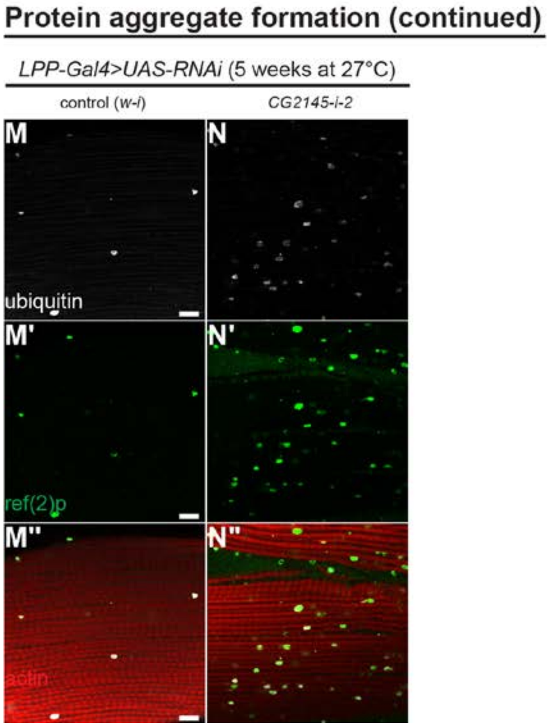


**Fig. S15: Adult flies with fat body (FB) RNAi (using *LPP-Gal4*) against *CG4332*, *CG2145*, and *CG31326* have increased muscle protein aggregate formation (Fig. 3 supplement).**

Shown are representative single confocal slices (cropped to equal extent for ease of visualization). Muscles were stained for ubiquitin, p62/ref(2)p, and actin (phalloidin). Quantification in **Fig. 3, D to H** was performed from these samples (using uncropped images). Scale bar: 10 µm.

**(A and B)** *LPP-Gal4>RNAi* (TRiP, Harvard Transgenic RNAi Project), 3 weeks old at 27°C.

**(C to F)** *LPP-Gal4>RNAi* (NIG, Japan National Institute of Genetics), 3 weeks old at 27°C.

**(G to I)** *LPP-Gal4>RNAi* (VDRC, Vienna *Drosophila* Resource Center), 3 weeks old at 29°C.

**(J to L)** *LPP-Gal4>RNAi* (NIG), 5 weeks old at 27°C.

**(M and N)** *LPP-Gal4>RNAi* (TRiP), 3 weeks old at 27°C.

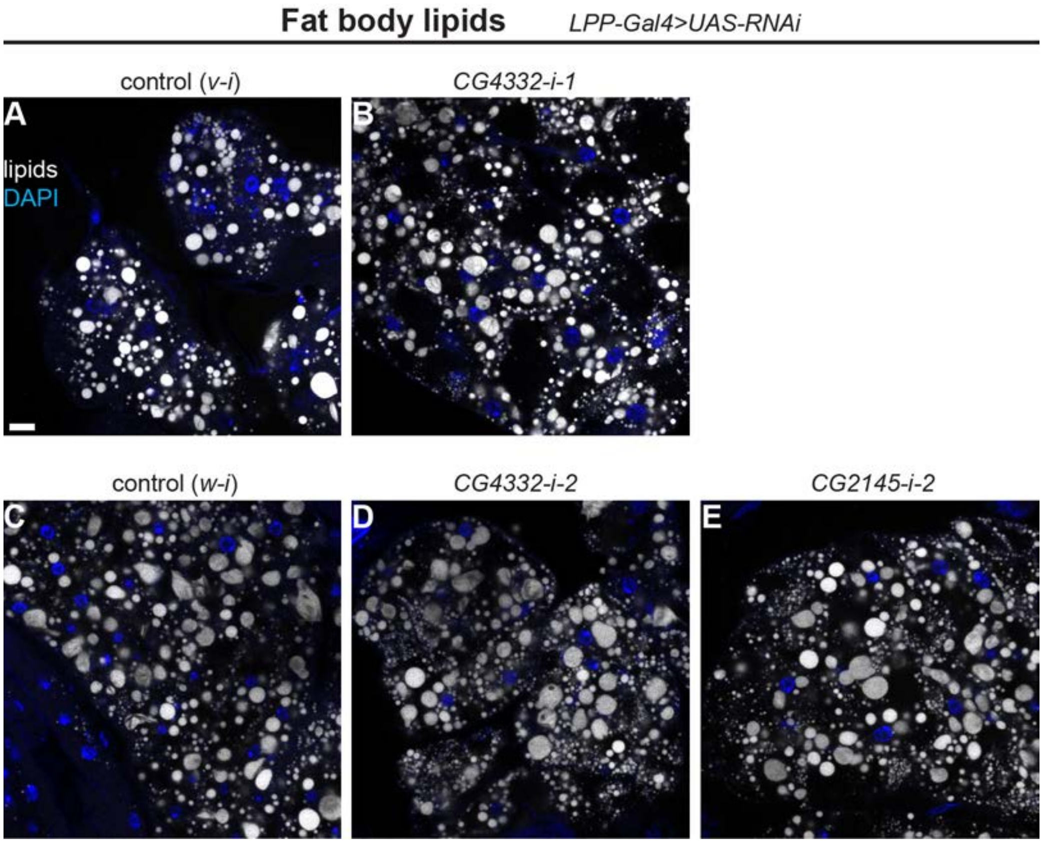


**Fig. S16: Adult flies with fat body (FB) RNAi (using *LPP-Gal4*) against *CG4332* or *CG2145* do not have significant defects in FB lipid droplets (Fig. 3 supplement).**

FBs were stained for lipids (BODIPY) and DAPI. Representative single confocal slices are shown. Scale bar: 10 µm.

**(A and B)** *LPP-Gal4>RNAi* (NIG, Japan National Institute of Genetics), 3 weeks old at 27°C.

**(C to E)** *LPP-Gal4>RNAi* (TRiP, Harvard Transgenic RNAi Project), 3 weeks old at 27°C.

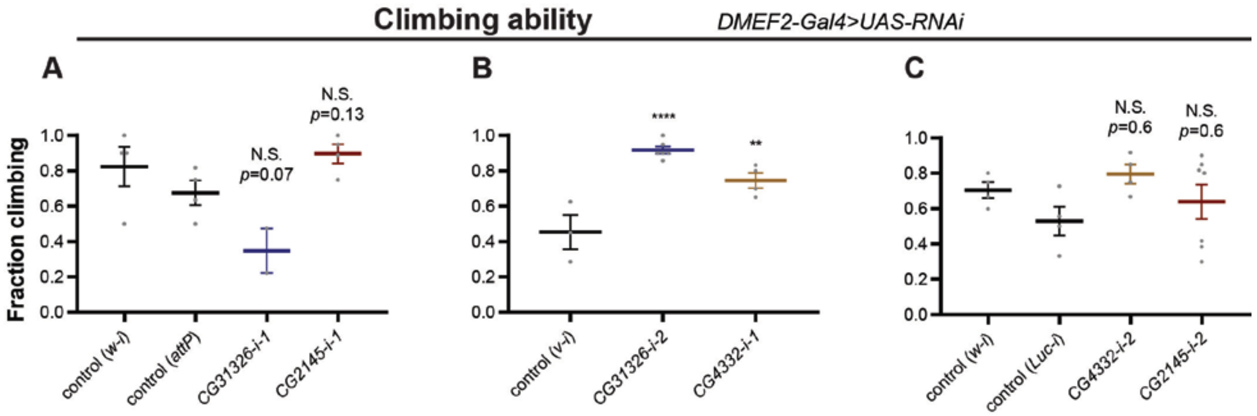


**Fig. S17: Adult flies with muscle RNAi (using *DMEF2-Gal4*) against *CG4332*, *CG2145*, or *CG31326* do not have significant defects in climbing ability (Fig. 3 supplement).**

**(A)** *DMEF2-Gal4>RNAi* from VDRC (Vienna *Drosophila* Resource Center) at 3 weeks old and 29°C. Biological replicates: *n*=4 (*w-i*), *n*=4 (*attP*), *n*=2 (*CG31326-i-1*), *n*=4 (*CG2145-i-1*). *P*-values are relative to control (*attP*).

**(B)** *DMEF2-Gal4>RNAi* from NIG (Japan National Institute of Genetics) at 5 weeks old and 27°C. Biological replicates: *n*=4 (*v-i*), *n*=6 (*CG31326-i-2*), *n*=4 (*CG4332-i-1*). *P*-values are relative to control (*v-i*).

**(C)** *DMEF2-Gal4>RNAi* from TRiP (Harvard Transgenic RNAi Project) at 5 weeks old and 27°C. Biological replicates: *n*=4 (*w-i*), *n*=4 (*Luc-i*), *n*=4 (*CG4332-i-2*), *n*=7 (*CG2145-i-2*). *P*-values are relative to control (*w-i*).

Statistics: mean±SEM; one-way ANOVA and Benjamini, Krieger, Yekutieli Linear Two-Stage Step-Up FDR.

**Table S1: Positive control secreted factors involved in intercellular signaling identified in total hemolymph MS**

| **Factor(s)** | **Fly hemolymph experiments** | **Human ortholog (DIOPT(*32*) and other references)** | **References** |
|---|---|---|---|
| IDGFs (imaginal disc growth factor) | most | CHIT1 | (*39, 80-85*) |
| ADGF-A (adenosine deaminase growth factor A) | N-glycan | CECR1 | (*86, 87*) |
| Tsp (thrombospondin) | N-glycan | THBS | (*88, 89*) |
| Ccn | N-glycan | CTGF | (*30, 90*) |
| spz (spätzle) | N-glycan | | (*91, 92*) |
| Sog (short gastrulation) | N-glycan | CHRD | (*93, 94*) |
| Nec (necrotic) | most | SERPINC1 | (*95*) |
| Spn5 (serpin 88Ea) | most | SERPINI1 | (*96, 97*) |
| Ance (Angiotensin converting enzyme) | most | ACE | (*98*) |
| Pvf3 (PDGF and VEGF related factor 3) | N-glycan | PDGF/VEGF | (*99*) |
| Upd1 (unpaired 1) | Neat, targeted | IL6 | (*62, 100*) |
| Egr (Eiger) | N-glycan | TNF-α | (*101*) |
| Tk (Tachykinin) | 3-10 kD | TAC1 | (*102*) |
| CCHa1 (CCH amide 1) | <3 kD, non-tryptic search | | (*103*) |
| SDR (secreted decoy of insulin receptor) | N-glycan | INSR | (*104*) |
| GBP2 (growth blocking peptide 2) | most | EGFs(*105*) | (*105*) |
| PGRPs (Peptidoglycan recognition proteins; multiple identified) | most | PGLYRPs | (*106*) |
| Mav (maverick) | neat (older flies) | TGF-β family | (*107*) |

**Note:** Upd1 was identified from a targeted MS analysis of unfractionated fly blood using a known tryptic peptide from ref.(*62*).

**Table S2: Identified positive control fat body to legs and muscle trafficking proteins**

| Factor | Function | Ref. | Human ortholog | DIOPT score(*32*) (version) | Dataset |
|---|---|---|---|---|---|
| Lsp1β (larval serum protein 1β) | Lsp complex consisting of α, β, γ subunits: FB-enriched expression; storage; cuticle contribution in several insect species; in *Calliphora vicina* contributes to muscle in thorax; protein abundance associated with adult wing size; Lsp1γ is important for and traffics to muscle adhesions. | (*108-113*) | | | BirA*G3-ER, BirA*R118G-ER |
| Lpp (apolipophorin) | Produced in FB; transport hydrophobic molecules including lipids, sterols, membrane lipids, and signaling secreted factors (Hh and Wg) to and in tissues including imaginal discs; interacts with lipophorin receptors. | (*7, 114-118*) | APOB (apolipoprotein B) | v.5: 4<br>v.6: 5 | BirA*G3-ER, BirA*R118G-ER |
| LTP (apolipoprotein lipid transfer particle) | Produced in FB and traffics to distal tissues including imaginal discs in which lipids get transferred from Lpp to imaginal disc cells through LTP; interacts with lipophorin receptors. | (*119, 120*) | APOB (apolipoprotein B) | v.5: 3<br>v.6: 4 | BirA*G3-ER |
| Idgf1 (Imaginal disc growth factor 1) | Secreted from FB and traffics to imaginal discs to promote cell proliferation; may act synergistically with insulin; may be regulated by miR-8 in FB and miR-8 may regulate whole-body growth; cuticle formation | (*39, 80, 81, 83, 84*) | CHIT1 (chitinase 1) | v.5: 1<br>v.6: 2 | BirA*G3-ER |
| Idgf2 (Imaginal disc growth factor 2) | Secreted from FB and traffics to imaginal discs to promote cell proliferation; may act synergistically with insulin; may be regulated by miR-8 in FB and miR-8 may regulate whole-body growth. | (*80, 81, 84, 85*) | CHIT1 (chitinase 1) | v.5: 1<br>v.6: 2 | BirA*G3-ER |
| Idgf6 (Imaginal disc growth factor 6) | Expressed in tissues including FB; may be regulated by miR-8 in FB and miR-8 may regulate whole-body growth; cuticle formation; interaction with chitin. | (*39, 80-83*) | CHIT1 (chitinase 1) | v.5: 1<br>v.6: 2 | BirA*G3-ER, BirA*R118G-ER |
| Adgf-D (adenosine deaminase-related growth factor D) | Secreted from FB and exerts proliferative effects on imaginal disc cells. Adgf-D mutant adults may be under-active. Adgf-D acts outside the cell and degrades extracellular | (*86, 87, 121-125*) | ADA2/CECR1 (adenosine deaminase 2 or cat eye syndrome chromosome | v.5: 7<br>v.6: 9 | BirA*G3-ER |

| | | | | | |
|---|---|---|---|---|---|
| | adenosine. Adenosine outside the cell has anti-proliferative, catabolic and energy store wasting effects, acting via adenosine transmembrane transport or adenosine receptor (AdoR), expressed in many tissues. AdoR or extracellular adenosine overexpression cause thoracic and wing defects, and AdoR overexpression causes defects in movement. | | region candidate 1) | | |
| cv-d (crossveinless d) | Secreted primarily from FB into hemolymph and traffics to imaginal discs. Binds to TGFs Gbb and Dpp and regulates their extracellular transport in imaginal discs. Gbb, Dpp, and cv-d also regulate adult muscle morphology. | (*7, 126, 127*) | | | BirA*G3-ER, BirA*R118G-ER |
| Tig (Tiggrin) | ECM protein secreted by FB (also blood cells, and to a lower extent possibly in muscles); binds to muscle adhesion foci through PS2 integrins and possibly other receptors; regulates muscle adhesion, thickness, and contraction velocity. | (*113, 128-130*) | TRIP11 | v.5: 1 | BirA*G3-ER |
| fon (fondue) | Produced by FB, secreted into hemolymph, and binds to muscle adhesion foci; regulates muscle adhesions and movement speed. | (*113*) | | | BirA*G3-ER |

**Abbreviations:** FB, fat body; Hh, hedgehog; Wg, wingless; TGF, transforming growth factor; Dpp, decapentaplegic; Gbb, glass bottom boat; ECM, extracellular matrix; Ref., references; v., version (DIOPT).

**Table S3: Identified additional known FB-produced proteins in *LPP-Gal4>BirA*G3-ER* legs**

| **Factor** | **Function** | **References** | **Human ortholog** | **DIOPT score(*32*) (version)** |
|---|---|---|---|---|
| Acer (Angiotensin-converting enzyme-related) | Peptidase produced by the FB and secreted to hemolymph; regulates sleep and heart functions. Mammalian ACE and ACE2 are expressed in adipose tissue and are involved in obesity. | (*131-136*) | ACE (angiotensin-converting enzyme), ACE2 (angiotensin-converting enzyme 2) | v.5: 6 (ACE)<br>v.6: 10 (ACE)<br>v.6: 6 (ACE2) |
| psh (Persephone) | Protease expressed in FB, upstream of SPE; within Toll pathway. | (*39, 137*) | TMPRSS12 | v.5: 1 |
| GNBP3 (Gram-negative bacteria binding protein 3) | Glucan recognition protein expressed in FB, upstream of SPE, within Toll pathway. | (*39, 137, 138*) | | |
| GNBP-like3 | Immune glucan-binding protein; may be produced by the FB. | (*39, 139*) | | |
| SPE (Spatzle Processing Enzyme) | Protease expressed in FB, cleaves pro-Spatzle to cause its activation. | (*39, 140, 141*) | | |
| TotA (Turandot A) | FB-produced peptide; expression changes due to stress. | (*39, 142-145*) | | |
| MP1 (Melanization Protease 1) | Protease expressed in FB; involved in melanization. | (*39, 146*) | | |
| Sp7 (Serine Protease 7) | Protease expressed in hemocytes and possibly in FB; involved in melanization. | (*39, 146, 147*) | F12, HGFAC | v.5: 1 |

**Abbreviations:** FB, fat body; v., version (DIOPT).

**Table S4: Additional secreted proteins identified in *LPP-Gal4>BirA*G3-ER* legs**

| **Factor** | **Function** | **References** | **Human ortholog** | **DIOPT score(*32*) (version)** |
|---|---|---|---|---|
| Fer1HCH (Ferritin 1 heavy chain homologue) | Ferritin iron carrier, may be secreted systemically. | (*148, 149*) | FTH1, FTHL17, FTMT | v.5: 1<br>v.6: 2 |
| Hmu (hemomucin) | A mucin with a putative signal peptide. | (*36, 150, 151*) | APMAP | v.5: 8<br>v.6: 8 |
| Ance-4, Ance-5 (Angiotensin-converting enzyme-4 and 5) | Expressed in FB, has a signal peptide related to Acer. | (*36, 39, 152*) | ACE | v.6: 3 |
| teq (tequila) | Protease expressed in FB and head; has a signal peptide; regulates lifespan, memory, and insulin signaling; mammalian PRSS12/neurotrypsin regulates social behavior and cognitive ability, and may be secreted. | (*36, 39, 153-158*) | PRSS12 (neurotrypsin) | v.5: 3<br>v.6: 2 |
| Nplp2 (Neuropeptide-like precursor 2) | Neuropeptide. | (*159-161*) | | |
| Tep2 (thioester containing protein 2) | Immune protein with a signal peptide; expressed in FB; *Anopheles gambiae* Tep1 is in the hemolymph. Mammalian CD109 may be secreted into the blood and may regulate TGF-β signaling. | (*36, 39, 162-168*) | CD109 | v.5: 7<br>v.6: 7 |
| tok (tolkin) or tlr (tolloid-related) | Secreted protease that processes pro-TGF secreted factors (e.g., dawdle) and inhibitors (e.g., short gastrulation); has functions in axon formation of motor neurons, and wing disc formation; expressed in FB. In *Drosophila* and mammals, TGF-β pathway regulates | (*39, 75, 93, 169-173*) | BMP1 (bone morphogenetic protein 1), TLL1 (tolloid like 1) | v.5: 7<br>v.6: 9 |

| | | | | |
|---|---|---|---|---|
| | metabolism, including muscles. In mammals, BMP1 and TLL1 proteases cleave and activate myostatin and GDF-11. | | | |
| modSP (modular serine protease) | Secreted protease in the Toll pathway. | (*174*) | several proteases including TMPRSS6, TMPRSS15, PAMR1, MASP2, MASP1, C1S, CSMD2 | v.5: 1<br>v.6: 1 |
| cathD (cathepsin D) | Protease with a signal peptide; loss of function causes neuronal defects. Mammalian secreted pro-CTSD has growth-factor activity, and CTSD proteolytically regulates growth factors and cytokines (e.g., insulin, glucagon, osteocalcin, IGFBP, FGF, interleukin-1, plasminogen); CTSD mutations result in neurodegeneration and psychomotor defects. | (*36, 175-182*) | CTSD (cathepsin D) | v.5: 9<br>v.6: 10 |
| Ndg (nidogen) | ECM protein. Mammalian protein is part of basement membrane. | (*183, 184*) | NID1 (nidogen 1) | v.5: 8<br>v.6: 10 |

**Abbreviations:** TGF, transforming growth factor; GDF, growth differentiation factor; IGFBP, insulin-like growth factor binding protein; FGF, fibroblast growth factor; ECM, extracellular matrix; v., version (DIOPT).

**Table S5: *Drosophila* orthologs of human secreted proteins involved in systemic pathways identified in BirA*G3-ER fat body to legs mass spectrometry.**

| **Mammalian factor(s)** | **Identified *Drosophila* ortholog with fly to human DIOPT score (v.6 or, if indicated, v.5)(*32*)** | **Mammalian function** | **References** |
|---|---|---|---|
| ACE (angiotensin-converting enzyme) | Acer (score=10), Ance-4 (score=3), Ance-5 (score=5) | Involved in obesity and expressed in adipose tissue. | (*131-136*) |
| APOB (apolipoprotein B) | Lpp (score=5), LTP (score=4) | Lipid transport between tissues; can bind to sonic hedgehog (SHH). Highly expressed in the liver. | (*7, 114-120, 185, 186*) |
| TLL1 (tolloid like 1), BMP1 (bone morphogenetic protein 1) | tok (score=9) | These factors cleave and activate myostatin and GDF-11, which are systemic TGF-β factors regulating metabolism, including muscles. | (*75, 93, 169-173, 187, 188*) |
| LNPEP (leucyl and cystinyl aminopeptidase) | CG4467 (score=4) | Circulates in blood during pregnancy. Cuts hormone peptides including oxytocin, met-enkephalin, vasopressin. May be involved in adipose tissue biology, thermogenesis, and glucose translocation. | (*189-195*) |
| TRHDE (thyrotropin-releasing hormone degrading enzyme), ENPEP (glutamyl aminopeptidase) | CG11951 (score=5 for LVRN, ANPEP; score=4 for TRHDE, ENPEP) | TRHDE: cleaves thyrotropin-releasing hormone. ENPEP: secreted during pregnancy; cuts cholecystokinin 8 and angiotensin 2. | (*196-199*) |
| ECE1 (Endothelin converting enzyme 1) | CG14526 (score=3), CG14528 (score=2), CG9507 (score=2) | Converts big endothelin to active endothelin via proteolysis. Endothelin in turn regulates organismal developmental progression, growth, disease progression, vascular constriction, and pressure of blood. ECE1 may be regulated by blood flow, metabolic dysfunction. ECE1 can be secreted through cleavage of the transmembrane form. | (*200-203*) |
| MME (membrane metalloendopeptidase) (also known as NEP/neprilysin) | CG14528 (score=2) | MME/NEP can be secreted through cleavage of the transmembrane form and detected in blood. Cuts hormone peptides including glucagon, somatostatin, insulin, substance P, cholecystokinin, enkephalins, angiotensins, bradykinin, natriuretic peptides, and others, thus playing roles in development, homeostasis, and disease. Expressed in the fat tissue along with other tissues. | (*186, 204, 205*) |
| KLKB1 (kallikrein B1) (also known as plasma | CG8586 (score=2) | KLKB1 or blood plasma-secreted kallikrein is part of the hormonal | (*186, 206-208*) |

| | | | |
|---|---|---|---|
| kallikrein) | | kallikrein-kinin (e.g., bradykinin) network. Kinins play roles in metabolism, diabetes, blood pressure control, inflammation. KLKB1 levels are related to metabolic disorders and diabetes. KLKB1 can also cleave plasminogen, and this regulates adipogenesis. Expression enriched in the liver. | |
| CD109 | Tep2 (score=7), Tep4 (score=5) | May be secreted into the blood and possibly regulate TGF-β signaling. | (*166-168*) |
| CTSD (Cathepsin D) | CathD (score=10) | CTSD regulates through cleavage hormones, cytokines and growth factors (such as IGFBP, FGF, glucagon, insulin, osteocalcin, interleukin-1, and plasminogen). Secreted pro-CTSD may be a growth factor. Also, phsychomotor and neurodegeneration defects may results from CTSD mutations. | (*177-182*) |

**Abbreviations:** TGF, transforming growth factor; GDF, growth differentiation factor; IGFBP, insulin-like growth factor binding protein; FGF, fibroblast growth factor; v., version (DIOPT).

**Table S6: Novel fat body to muscle communication factors suggested by our data**

| Factor | Putative human orthologs (with references) | DIOPT score (version) (*32*) | SP (*36*) | *LPP-Gal4>RNAi* muscle phenotype (2 lines) | *LPP-Gal4>RNAi* FB phenotype | *DMEF2-Gal4>RNAi* muscle phenotype | Dataset | MS score |
|---|---|---|---|---|---|---|---|---|
| CG4332 | CLPTM1L (cisplatin resistance related protein or cleft-lip and palate-associated transmembrane protein 1-like) – no known functions in adipose or muscle tissues. Variations in DNA sequence and expression levels are associated with tumor division and apoptosis (*209-212*). | v.6: 10 | Y | climbing ability↓ protein aggregates↑ | No significant effects on FB lipid-droplets | No consistent, statistically-significant climbing-ability defects | BirA*R118G-ER | 2 |
| CG2145 | ENDOU (poly-U-specific placental endonuclease) – signal-peptide-containing RNA-binding protein of uncharacterized function in muscles or adipose tissue; has a role in B lymphocytes (*213-215*). | v.5: 7<br>v.6: 9 | Y | climbing ability↓ protein aggregates↑ | No significant effects on FB lipid-droplets | No consistent, statistically-significant climbing-ability defects | BirA*G3-ER | 4 |
| CG31326 | 1. PAMR1 (peptidase domain-containing associated with muscle regeneration-1) – expression associated with muscle regeneration (*216*) (but no functional data in muscles or adipose tissue), and overexpression in breast cancer cells may reduce growth *in vitro* (*217*)<br>2. Coagulation factor FVII – may be secreted by adipose tissue and liver and is associated with obesity, insulin resistance, type-2 diabetes, and high fat diet feeding (*218, 219*). | 1. v.5: 1 (PAMR1)<br>2. v.6: 1 (FVII) | Y | climbing ability↓ protein aggregates↑ | | No consistent, statistically-significant climbing-ability defects | BirA*G3-ER | 4 |

**Abbreviations:** SP, signal peptide; FB, fat body; MS, mass spectrometry; v., version (DIOPT)
